# Population genomics of ancient and modern *Trichuris trichiura*

**DOI:** 10.1101/2021.10.21.464505

**Authors:** Stephen R. Doyle, Martin Jensen Søe, Peter Nejsum, Martha Betson, Philip J. Cooper, Lifei Peng, Xing-Quan Zhu, Ana Sanchez, Gabriela Matamoros, Gustavo Adolfo Fontecha Sandoval, Cristina Cutillas, Louis-Albert Tchuem Tchuenté, Zeleke Mekonnen, Shaali M. Ame, Harriet Namwanje, Bruno Levecke, Matthew Berriman, Brian Lund Fredensborg, Christian Moliin Outzen Kapel

**Affiliations:** Wellcome Sanger Institute, Hinxton, Cambridgeshire, CB10 1SA, United Kingdom; Department of Plant and Environmental Sciences, University of Copenhagen, 1871 Frederiksberg, Denmark; Department of Clinical Medicine, Aarhus University, 8220 Aarhus N, Denmark; School of Veterinary Medicine, University of Surrey, Daphne Jackson Road, Guildford, GU2 7AL, United Kingdom; Institute of Infection and Immunity, St George’s University of London, London, SW17 ORE, United Kingdom; School of Medicine, Universidad Internacional del Ecuador, Quito, Ecuador; Department of Parasitology, School of Basic Medical Sciences, Guangdong Medical University, Zhanjiang, Guangdong Province 524005, People’s Republic of China; College of Veterinary Medicine, Shanxi Agricultural University, Taigu, Shanxi Province 030801, People’s Republic of China; Department of Health Sciences, Brock University, St. Catharines, Ontario, L2S 3A1, Canada; Microbiology Research Institute, Ciudad Universitaria, Universidad Nacional Autónoma de Honduras, Tegucigalpa, Honduras; Departamento de Microbiología y Parasitología, Facultad de Farmacia, Universidad de Sevilla, Sevilla, Spain; Department of Translational Physiology, Infectiology and Public Health, Ghent University, Ghent, Belgium; Vector Control Division, Ministry of Health, P.O.Box 1661, Kampala, Uganda; Faculty of Sciences, University of Yaoundé I, P.O. 7244 Yaoundé, Cameroon; Institute of Health, School of Medical Laboratory Sciences, Jimma University, Jimma, Ethiopia; Public Health Laboratory Ivo de Carneri, Pemba, Tanzania

## Abstract

The neglected tropical disease trichuriasis is caused by the whipworm *Trichuris trichiura*, a soil-transmitted helminth that has infected humans for millennia. Today, *T. trichiura* infects as many as 500 million people, predominantly in communities with poor sanitary infrastructure enabling sustained faecal-oral transmission. Using whole-genome sequencing of geographically distributed worms collected from human and other primate hosts, together with ancient samples preserved in archaeologically-defined latrines and deposits dated up to one thousand years old, we present the first population genomics study of *T. trichiura*. We describe the continent-scale genetic structure between whipworms infecting humans and baboons relative to those infecting other primates. Admixture and population demographic analyses support a stepwise distribution of genetic variation that is highest in Uganda, consistent with an African origin and subsequent translocation with human migration. Finally, genome-wide analyses between human samples and between human and non-human primate samples reveal local regions of genetic differentiation between geographically distinct populations. These data provide insight into zoonotic reservoirs of human-infective *T. trichiura* and will support future efforts toward the implementation of genomic epidemiology of this globally important helminth.

## Introduction

The human-infective whipworm, *Trichuris trichiura*, is a soil-transmitted helminth (STH) responsible for trichuriasis, a neglected tropical disease (NTD) estimated to affect as many as 500 million people worldwide ^1^. As an intestinal parasite, an infection begins by the ingestion of embryonated eggs in contaminated food or soil; eggs migrate to the large intestine and hatch, after which emerging larvae burrow and establish an intracellular niche within intestinal epithelia ^2^ where they develop to adult stages that can remain *in situ* for years ^3^. Although low and moderate infection burdens typically remain asymptomatic, favouring efficient transmission in regions with poor sanitation such as endemic rural settings, chronic infections with high worm burdens cause a range of debilitating gastrointestinal symptoms and can lead to nutritional deficiencies and delays in physical and cognitive development, especially in children ^4^. Infections with *T. trichiura* (and the other STHs, i.e. *Ascaris lumbricoides* and the hookworms *Ancylostoma duodenale, A. ceylanicum*, and *Necator americanus*) are primarily treated with benzimidazole anthelmintics (albendazole or mebendazole), although the efficacy of these drugs for the treatment of *Trichuris* is suboptimal ^5,6^ and few alternative drugs are available; in endemic regions, treatment is targeted towards pre-school and school-aged children, women of reproductive age as well as high-risk working adults via annual or biannual mass drug administration (MDA) campaigns. In 2019, more than 613.1 million children, representing 60% treatment coverage of children at-risk of STH infection, were treated with anthelmintics ^7^. It is almost certain that this number will only increase in the near future, since STH are targeted for elimination as a public health problem by 2030, in line with the WHO Sustainable Development Goals for NTDs ^8^ and the WHO road map for NTDs 2021—2030.

Humans have been parasitised by *T. trichiura* for millennia. Although now generally restricted to tropical and subtropical regions ^1^, this parasite was once a globally distributed worm. Parasite eggs have been found in human coprolites (fossilised faeces) from archaeological sites dated back to 7,100 BC ^9–11^, including sites in Europe and North America where autochthonous infections are now unusual ^12–16^. However, whipworms are known to infect a broad range of mammals; over 70 species have been described within the genus *Trichuris*, and while generally host-specific, cross-host species transmission of individual parasite species have been reported, including between humans and pigs ^17^, humans and dogs ^18^, or humans and non-human primates ^19–23^. Parasites that can interchangeably infect human and non-human hosts represent a credible challenge to control campaigns, as non-human hosts may act as a reservoir in which parasites can evade treatment and subsequently become a source of infections for treated populations. Understanding the historical and modern dispersal of *T. trichiura* in human and non-human hosts is likely required to achieve elimination. Further, this goal of elimination of STH-attributable morbidity by MDA is threatened by the emergence of resistance to the benzimidazole compounds used to control them, as has been observed in other parasitic worms (in particular, veterinary helminths) frequently exposed to the drug ^24^. Although evidence to suggest that resistance is emerging in *T. trichiura* is limited, understanding population connectivity within and between endemic regions will inform the likelihood and rate by which resistance alleles might spread when they arise.

Here, we describe the first population genomic analysis of *T. trichiura*. Using whole-genome sequencing data of modern whipworms collected from human and non-human primate hosts, together with ancient samples preserved in archaeological defined latrine and deposits ^13^, we describe broad- and fine-scale genetic structure, admixture, and population demographics of geographically and genetically distinct populations. We also explore genome-wide evidence of local adaptation between different human worms and between human and animal worms. These genome-wide variant data will support future efforts toward the implementation of genomic epidemiology of this globally important STH.

## Results

### Sequencing of ancient and modern *Trichuris trichiura*

We have generated whole-genome sequencing data from 44 modern and 17 ancient samples (**Supplementary Table 1**), resulting in an average coverage of 9.31× and 0.66× of the nuclear genomes and 613× and 95× of the mitochondrial genomes, respectively (**Supplementary Table 2**). The samples analysed were derived from a broad geographic distribution and include 18 populations from nine countries spanning Africa, Central America, Asia, and Europe (**Fig. 1a**). The modern samples were composed of 37 worms obtained from human hosts, as well as seven samples obtained from captive animal hosts, including two parasites from baboons (*Papio hamadryas*), two from colobus monkeys (*Colobus guereza kikuyensis*) and three from leaf monkeys (*Trachypithecus francoisi*).

**Figure 1.**
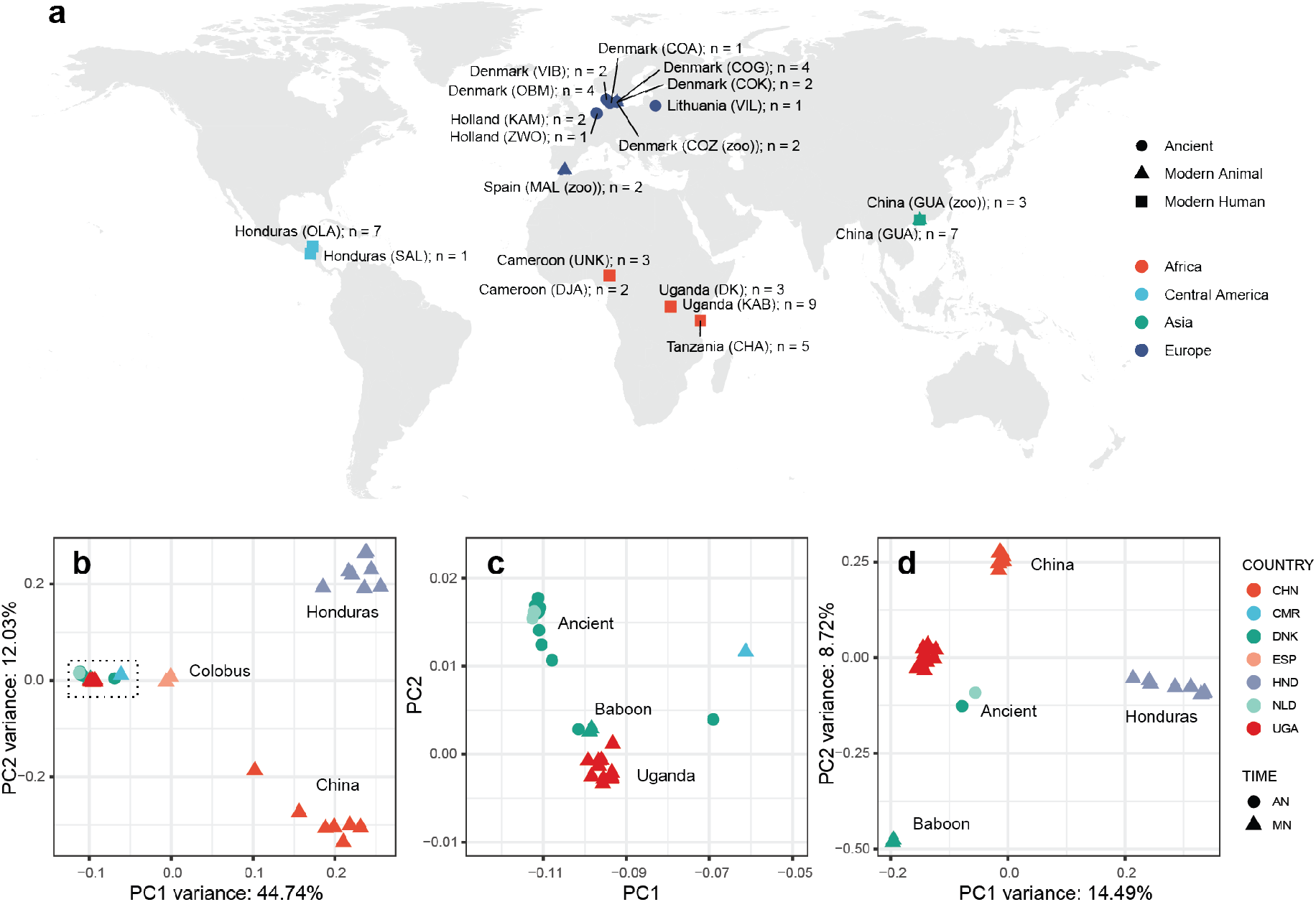
Global sampling distribution and broad-scale genetic relatedness of ancient and modern *Trichuris trichiura*. **a**. World map showing the approximate sampling locations of samples used in the study, highlighting geographic regions of ancient and modern sampling, as well as host species. **b**. Principal component analysis of mitochondrial diversity (56 samples, 802 variants). **c**. Zoomed-in view of the cluster of samples indicated by the dashed box in panel b, containing ancient, Ugandan, and baboon-derived samples. **d**. Principal component analysis of nuclear diversity using a subset of higher quality samples (31 samples, 2,631,365 variants). Sample abbreviations: CHN=China; CMR=Cameroon; DNK=Denmark; ESP=Spain; NLD=Netherlands; UGA=Uganda; HND=Honduras; AN=ancient; MN=modern.

The ancient samples were obtained from archaeological latrines and dig sites, primarily from Denmark (five locations) as well as from the Netherlands (two locations) and a single site in Lithuania, which individually have been dated to span the last thousand years (**Supplementary Fig. 1**). Thus, these samples represent the oldest helminth samples, and likely the oldest eukaryotic pathogens ^25^, from which whole-genome sequencing data has been derived to date. Reads derived from ancient samples displayed increased sample damage due to deamination relative to the modern samples (**Supplementary Fig. 2; Supplementary Table 2**), particularly in the first two bases of each read, which were removed before downstream analyses.

Joint genotyping followed by stringent filtering identified a total of 1,888 mitochondrial and 6,933,531 nuclear variants in the sample cohort. Considering the variation in depth of coverage and degree of variant missingness between samples, we separately analysed subsets of samples with either mitochondrial or nuclear sequence data using variants with at least 3x coverage across at least 80% of sites within each dataset. Depth of coverage analyses comparing autosomal and sex-linked scaffolds revealed 17 male (XY; expected 0.5× depth of sex-linked scaffolds relative to autosomes) and 17 female (XX; expected 1:1 ratio of sex-to-autosomal depth) worms in the individual worm sequencing data; the datasets derived from pooled eggs (ancient samples, modern samples from Cameroon & Tanzania) revealed intermediate coverage due to the presence of mixed-sex within the pools (**Supplementary Fig. 3; Supplementary Table 2**). Depending on the analysis, the colobus and leaf monkey samples were also excluded.

### Broad-scale genetic diversity of ancient and modern helminths

To investigate the broad-scale genetic diversity within the global cohort, we first performed a principal component analysis of mitochondrial variation (**Fig. 1b**). Three genetically defined clusters were identified, clearly separating samples from China and Honduras from the third cluster containing samples of mixed origin. Closer inspection of this third cluster identified closely related but genetically distinguishable ancient, modern Ugandan and baboon samples (**Fig. 1c)**. To provide further granularity, we analysed nuclear genetic variation with a subset of higher quality samples, which provided more apparent differentiation between geographically distinct human worms, as well as between the ancient and baboon samples (**Fig. 1d**). We also sampled additional populations from Africa (Cameroon & Tanzania), however, there was limited genetic information due to the very low sequencing coverage obtained, likely due to the fact we sampled only unembryonated eggs (with environmentally robust eggs shells, and thus, difficult to crack) rather than adult worms, and thus, the DNA concentration was very low and suboptimal for sequencing ^26^. In an attempt to address this, we reassessed genetic relatedness using identity-by-state (IBS) and IBS covariance analyses using ANGSD, which is more tolerant to low-coverage and sparse genetic data ^27^; here, a low but distinguishable genetic signal based on within-Africa clustering of these samples was identified (**Supplementary Fig. 4 & 5**). However, further sampling would be needed to precisely place the samples from these regions relative to the larger and better genetically defined cohort.

Whipworms obtained from leaf and colobus monkeys were highly genetically distinct from the human/baboon group and from each other (**Supplementary Fig. 6a**). To better understand their phylogenetic placement in the *Trichuris* genus, we assembled mitochondrial genomes from all modern worm samples using an iterative baiting and mapping approach and compared them to publicly available whole mitochondrial genomes. Whipworms from leaf monkeys formed a sister group to the human and non-human primate clades, whereas those from the colobus group were more closely aligned with pig whipworm *T. suis* samples in a separate cluster (**Supplementary Fig. 6b**). This finding provides further support that these animal-derived *Trichuris* spp. are genetically distinct species from the human- and baboon-infective *T. trichiura*.

### Patterns of admixture between populations

To more formally describe the genetic relationships between populations, we first used NGSadmix to visualise the ancestral composition of samples. We estimated three ancestry components (K = 3) (**Fig. 2a**), broadly consistent with the global population structure in mitochondrial and nuclear PCAs (**Fig. 1b,c,d**); ancient and baboon samples formed a cluster that was distinct from each of China and Honduras populations. Ugandan samples showed mixed ancestry of China and ancient/baboon samples. To explore this further, we determined admixture proportions across a range of K values (K = 2–10) (**Supplementary Fig. 7**). Subtle evidence of shared ancestry between China and Honduras was observed, but this disappears at K ≥ 3. Ugandan samples displayed the most diverse ancestry profile, with complex mixtures of private components not found elsewhere; we note that the first two Ugandan samples of the admixture plot that maintain the same admixture profile throughout were sampled from the same host and are likely 2^nd^ degree relatives (MN_UGA_DK_HS_001 & MN_UGA_DK_HS_002; kinship coefficient [Θ] = 0.34). Similarly, samples from Honduras also increased in diverse components throughout the range. Interestingly, the ancient and baboon samples maintained a single fixed ancestry profile throughout the conditions until K = 10, where a single baboon sample diverged.

**Figure 2.**
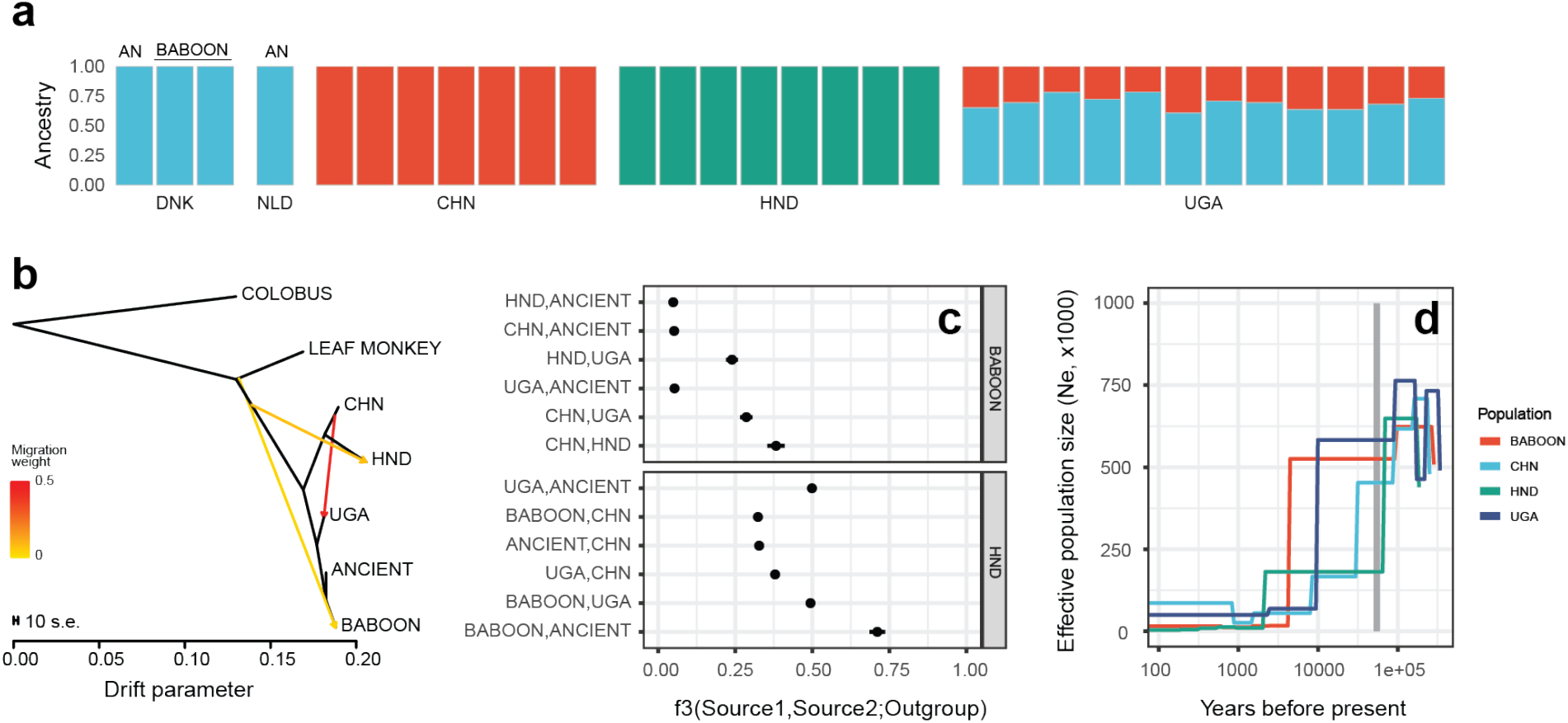
Fine-scale genetic relationships and admixture between global populations. **a**. Admixture plot depicting population ancestry proportions determined using NGSadmix (K = 3, variants = 1,001,427). See Supplementary Fig. 7 for a complete analysis of K from 2 to 10. Ancient (AN) and baboon samples are highlighted. **b**. Treemix maximum likelihood tree of ancient and modern samples, including Colobus and Leaf monkey samples as outgroups, showing two migration edges. See Supplementary Fig. 8 for a complete analysis of migration edges from 0 to 5. **c**. Outgroup *f*3 statistics were determined using *qp3Pop* in ADMIXTOOLS to compare allele frequency correlations between two source populations (indicated on the left side of the panel), relative to an outgroup population (right side of the panel). Positive *f*3 values imply shared or correlated genetic drift between Source 1 and Source 2 populations. Whiskers represent the standard error, calculated using a weighted block jackknife. For all data presented, the Z score was significant (|Z| > 3). **d**. Population demographic history of each population was determined using SMC++ to compare effective population size (*N*e) over recent evolutionary history. The grey vertical box highlights the period between 50—60 thousand years ago, coinciding with the migration of modern humans out of Africa. Sample abbreviations: CHN=China; UGA=Uganda; HND=Honduras.

To begin to quantify admixture, we used Treemix to estimate migration between nodes under a maximum likelihood framework (**Fig. 2b**). Three migration edges were supported (see **Supplementary Fig. 8** for trees and associated residual heatmaps for edges = 0-5), and emphasised admixture between Ugandan and Chinese populations. Interestingly, there was evidence implicating a lower level of admixture between the base of the tree adjacent to the leaf monkey branch and both Honduran and Baboon samples. While this does not directly implicate hybridisation between non-human-infective and human-infective whipworms specifically, hybridisation between the phylogenetically related but pig-infective *T. suis* and *T. trichiura* has been described previously in Ecuador ^28^. We further quantified these admixture results using *f*-statistics on the nuclear variants (**Fig. 2c**). Using baboon samples as an outgroup, we further identified the relationship between China and the Honduras populations, and found that this relationship is closer than between Ugandan and Honduras populations. Investigations of mitochondrial markers of *rrn*L and *nad*1 from humans in Uganda, China and Ecuador (geographically close to Honduras) have identified close genetic similarity between parasites from China and Ecuador, and that these parasites are distinctly different to those from Uganda ^17^. Using baboon as an outgroup reduced our ability to measure more subtle differences between the closely related baboon, ancient, and Ugandan populations; to address this, we used Honduras as an outgroup, which provided a greater resolution to show the closer relationship between the baboon and ancient samples, relative to comparison with the Ugandan samples.

Finally, we used population demographic analyses to characterise the effective population size of the parasite populations over time (**Fig. 2d**), revealing that current populations are a fraction of the size of the historical populations from which they were derived. All populations showed a substantial decline in effective population size between 50 and 100 thousand years ago; Ugandan and baboon populations did maintain a significantly higher population size until less than ten thousand years ago, after which they underwent further decline. These findings, together with admixture analyses, allow us to hypothesise about the potential dispersal of *T. trichiura*. Although we have made specific assumptions about important but unmeasurable parameters of the population demographic model — specifically, mutation rate and generation time — and therefore the precision of these estimates is low, these data are consistent with bottlenecks in parasite populations that likely occurred as modern humans migrated out of Africa approximately 50 to 60 thousand years ago ^29^. The intermediate *f*3 statistics between Uganda and China suggests a stepwise migration pattern, first from Africa to Asia and then from Asia to the Americas, the latter of which would be consistent with human migration between the two continents, perhaps via a crossing of the Beringia land bridge. This migration was distinct to that which was established in Europe, indicated by the lower degree of shared variation between ancient and Chinese samples, but almost no sharing between the ancient and the Honduran samples. It has been argued that some intestinal parasites may not have survived the Beringia crossing due to unfavourable environmental conditions to sustain their lifecycle, and thus, genetic connectivity in parasite populations reflect a separate peopling of the Americas through trans-pacific or coastal migrations ^9^. To explore this here, we compared the proportion of variation shared between Ugandan, Chinese and Honduran samples. While a large proportion of variation was private (in total, 46.6% of variants are present only in one of the three populations) or shared by all three populations (26.8%), 3.2% of variants are shared between the Ugandan and Honduran populations that are absent from Chinese samples (**Supplementary Fig. 9**); while tenuous due to the low sample numbers and sparse geographic sampling, such variation would support a low level but independent introduction of parasites to the Americas from Africa.

### Genome-wide patterns of selection and adaptation

Distinct geographic clustering of Ugandan, Chinese, and Honduran populations prompted us to examine evidence of diversifying selection that might indicate local adaptation within the human-infective parasites. Mean nucleotide diversity (*π*) varied between populations, with the highest diversity observed in Uganda (genome-wide mean *π* = 0.0072), followed by China (*π* = 0.0058), baboon (*π* = 0.0051), and then Honduras (*π* = 0.0041), consistent with a global dispersal of *T. trichiura* originating in Africa (**Supplementary Fig. 10**). Ancient samples presented with the lowest diversity of all populations (*π* = 0.0013). Tajima’s D was consistent, on average, between modern human populations (**Supplementary Fig. 11a**), however, discrete regions of positive and negative values were observed genome-wide **(Supplementary Fig. 11b)**. Although all populations were on average above zero genome-wide, the ancient samples showed a distinct positive skew in Tajima’s D, consistent with a contracting population. The baboon population presented with a bimodal distribution, as a result of high levels of positive Tajima’s *D* on the sex-linked scaffolds; this observation is likely explained by the fact that both worm samples were male (**Supplementary Fig. 3**; MN_DNK_COZ_PH_001 & MN_DNK_COZ_PH_002) and thus, lacked variation as a result of being hemizygous. A genome-wide analysis of divergence, calculated using a sliding window of *F*_ST_ between pairs of populations, identified local regions of differentiation that may be indicative of local adaptation (**Fig. 3a**). For each pairwise comparison (UGA vs CHN: Weir and Cockerham weighted *F*_ST_ = 0.137; UGA vs HND: *F*_ST_ = 0.287; CHN vs HND: *F*_ST_ = 0.296), we identified 58, 49 and 53 genes in region of high divergence (defined as the top 1%), respectively. Analysis of gene ontology (GO) terms for each of the genesets (**Supplementary Tables 3, 4, and 5** for each pairwise comparison, respectively) did not, however, identify any enriched terms that would have suggested selection of multiple genes of common function. We extended this analysis to compare Ugandan and baboon samples, given their close genetic relationship despite being isolated from different host species. Genome-wide mean *F*_ST_ was intermediate (*F*_ST_ = 0.179) to that of the human-specific analyses, re-emphasising that despite divergence, baboon-infective whipworms are within the species-level of diversity expected of human-infective *T. trichiura*. Genome-wide, discrete regions of differentiation were observed, particularly on the sex-linked scaffolds (**Fig. 3b**); regions of high divergence (top 1%) contained 61 genes (**Supplementary Table 6**), however, as in the previous comparison, no GO term enrichment was observed.

**Figure 3.**
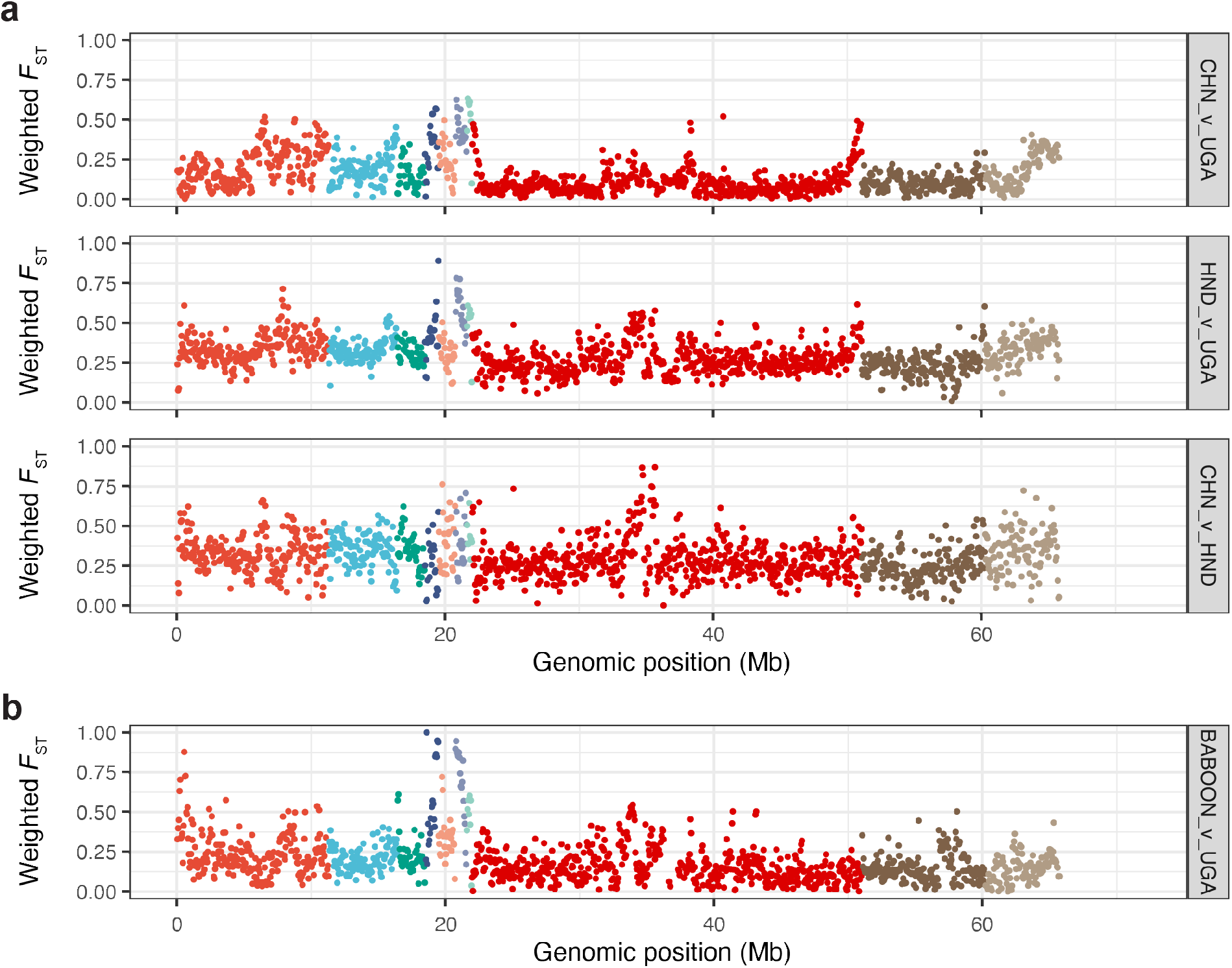
Genome-wide comparison of genetic variation. **a**. Comparison of human-infective *T. trichiura* from genetically and geographically defined populations. Pairwise *F*_ST_ was measured in 50 kb windows between China and Uganda (top), Uganda and Honduras (middle), and China and Honduras (bottom). **b**. Comparison of closely related human-infective Ugandan and baboon-infective *T. trichiura*. Sample abbreviations: CHN=China; UGA=Uganda; HND=Honduras.

Anthelmintic resistance could have catastrophic consequences were it to emerge in human infectious parasites. It was, therefore, important to use this global collection to explore variation in and around β-tubulin, a gene long associated with variation in response to benzimidazole-class anthelmintics such as those used to treat trichuriasis. We identified nine polymorphic sites in the sex-linked β-tubulin gene (Gene id: TTRE_0000877201; Location: Trichuris_trichiura_1_001: 10684531..10686350) segregating within modern human populations, however, no variation within the codon positions P167, P198, or P200 that are typically associated with benzimidazole resistance was observed (**Supplementary Fig. 12a**), consistent with previous studies ^23,30–32^. Further, there was little evidence of broader-scale genetic change on standing genetic variation in the region surrounding the β-tubulin gene that might be associated with positive selection on a gene within that region (**Supplementary Fig. 12b**,**c**).

## Discussion

The whipworm *T. trichiura* is one of the most prevalent and globally distributed helminth pathogens of humans. Here, we have undertaken a broad survey of genome-wide genetic diversity of *Trichuris* collected from human and non-human primates, including, to our knowledge, the oldest eukaryotic pathogens analysed using whole-genome sequencing to date. *Trichuris* spp., as a soil-transmitted helminth, completes part of its life cycle in the environment; unembryonated eggs are deposited in faeces where they undergo embryonation over two-to-four weeks under ideal, typically warm and moist, conditions commonly found in tropical and sub-tropical regions. The remarkable resilience of eggs in the environment allows them to remain viable in the soil for at least a decade ^33^ and morphologically identifiable to genus level after thousands of years ^9–11^. Here we demonstrate that DNA within eggs remains sufficiently preserved for whole-genome sequencing from samples up to one thousand years old, providing a unique and information-rich insight into parasites of the human past.

Analyses of the genomic diversity of ancient parasites and their genetic connectivity to modern parasite populations unexpectedly revealed a close genetic relationship to Ugandan and Baboon samples. Unfortunately, the ancient samples consisted of pools of eggs rather than individual worms, limiting investigation of population demographics and estimated split times between ancient and these closely related modern populations. Epidemiological estimates of *T. trichiura* prevalence have been proposed to be comparable between medieval populations in Europe (dated between 680 and 1700 CE) and modern, non-European endemic regions ^16^, and in Denmark, *Trichuris* infections were prevalent in children particularly from rural areas up to at least the 1930s ^34^. In contrast, the low genome-wide genetic diversity and positive Tajima’s D of our ancient samples are consistent with a constrained and likely contracting population size. Whether this reflects true population decline at the time the eggs were deposited, non-random sampling of genetically related parasite eggs within a defined archaeological site, or non-random distribution of whipworm prevalence in the host population is unclear. However, it is likely that the population demographics of these ancient northern European populations are different with respect to populations found in more tropical conditions where it is currently endemic. The rate of parasite development in the environment is temperature-dependent and decreases in cooler temperatures ^35^, and below 15°C, development arrests. This is particularly relevant for parasite populations such as those once endemic in northern Europe, as the proportion of the year during which favourable conditions for parasite development and transmission decreases at increasing latitudes. Thus, suboptimal environmental conditions necessary for parasite development and infection may in fact explain in part regional differences in parasite prevalence including local regions of population decline and diversity. Genomic analysis of ancient parasites from southern Europe and Africa will likely shed light on the missing links connecting the northern European and African parasites analysed here and the relationship between genetic diversity and endemicity in response to a changing climate.

Our genomic analysis of modern human and animal parasites provides further support that *T. trichiura* is specific to humans, baboons and maybe some other non-sampled primates and emphasises that parasite demography will be dependent on that of its host. Population demographic analyses of human infective whipworms support the hypothesis that global parasite populations suffered severe population bottlenecks coinciding with human migration out of Africa and that differential proportions of genetic admixture between these populations provide a plausible hypothesis by which parasites continued to infect modern humans as they distributed throughout the world. We note that one potential source of independent introduction of *T. trichiura* from Africa to the Americas resulting in shared parasite ancestry would likely be associated with the mass forced migration of Africans during the trans-Atlantic slave trade. We hypothesise that this genetic signature would be greater between West African (for which we have no samples) and American parasites than between Central or East African parasites as sampled here, and that West African parasites would be more suitable to test the degree of genetic admixture and thus impact of different migration events on the genetic diversity of endemic American parasites present today. Collectively, our data reinforce the important role that human migration has played on the spread of modern helminths throughout the world ^9,17,36–38^.

Comparative analysis of human and animal infective whipworms demonstrate that worms isolated from baboons likely represent a zoonotic reservoir of human-infective parasites in Africa. *T. trichiura* is a soil-transmitted pathogen and, unlike many zoonotic transmissions that involve a vector species, the likelihood of cross-species zoonotic transmission will be influenced by the degree to which humans and animals occupy and directly share ecological niches, for example, ground or water supply. Baboons are large ground-dwelling primates that can be found close to human activity and likely share a common source of infection of STHs such as *T. trichiura*. This is in contrast to smaller, tree-dwelling non-human primates including leaf and colobus monkeys, that would have less interaction with humans. However, genome-wide analyses between human and baboon parasites do reveal discrete regions of the genome that differ between hosts, potentially indicative of some degree of host adaptation. We note that whipworms were collected from only three species of captive non-human primates and we lack sampling data from other apes that live in proximity to humans, particularly in natural settings. Further sampling of human- and a broader range of animal-infective parasites, particularly where cohabitation exists, combined with genome-wide analyses of selection, should provide further insight into the mechanisms by which host-specificity is maintained, and cross-species transmission is tolerated.

Large-scale treatment of parasites using benzimidazole anthelmintics (as used by MDA programmes against STH including *T. trichiura*) is known to select genetic variation in β-tubulin conferring resistance to treatment. This is particularly evident in veterinary helminths ^36,39^, where selection for benzimidazole resistance has rendered benzimidazole derivatives almost ineffective in some regions of the world, particularly against helminths of horse and sheep ^24^. Emerging evidence suggests that MDA may be exerting similar selection pressure on STHs ^40–42^. We found no evidence for selection within or surrounding the β-tubulin gene. However, we do not have any data on their phenotypic response or exposure history to anthelmintic treatment, and thus, we cannot determine whether the absence of known benzimidazole-resistance associated variants in our data indicates that (i) all populations are phenotypically susceptible to anthelmintic treatment, or (ii) populations may or may not be phenotypically susceptible to benzimidazole treatment, but that selection on variation elsewhere in the genome, and not on variation in the β-tubulin gene, is responsible for drug response in *T. trichiura*. Future studies exploiting genome-wide analyses on parasite populations that are phenotypically well-defined for drug response ^43^ are needed to define the genetic architecture of drug response in *T. trichiura*. It will be particularly important to disentangle known variation in the tolerance to benzimidazole drugs by *T. trichiura* ^*44,45*^ from the emergence of loss of efficacy due to high drug exposure ^6^, especially if the promise of using molecular diagnostics to monitor the emergence of anthelmintic resistance is to be realised ^46–49^.

Our data establish the genetic framework for the genomic epidemiology of *T. trichiura*. It is almost certain that genomic surveillance will become an important tool for informing helminth control campaigns in the future ^50,51^. Such data could be used to interpret variation between and changes within parasite populations, for example, to characterise population decline due to effective control or distinguishing between transmission, recrudescence, and loss of efficacy of the drugs used to control them. These require a comprehensive understanding of the underlying genetic diversity of the parasite throughout its range. Our data provide a significant first step towards this goal and will be enriched by further sampling throughout endemic regions to characterise finer-scale genetic connectivity and effective parasite transmission zones.

## Acknowledgements

We gratefully acknowledge: Eske Willerslev, Kurt H. Kjær, and Martin Sikora for helpful discussions during the early stages of the project; Inga Merkyte, Mette Marie Hald, Kirsten Haase, Rikke Simonsen and Ruben Habraken for access to ancient samples; and Mads Frost Bertelsen for providing samples from baboons.

This work was funded by the University of Copenhagen’s KU2016 initiative, ‘The Genomic History of Denmark’, a UKRI Future Leaders Fellowship [MR/T020733/1], and the Wellcome Trust [206194]. For the purpose of Open Access, the author has applied a CC BY public copyright licence to any Author Accepted Manuscript version arising from this submission.

## Author contributions

CMOK, PN, and MJS designed the study. MJS extracted DNA from ancient samples and prepared NGS libraries for sequencing for all samples. PN extracted DNA from modern worm samples. MBet, PJC, LP, XQZ, AS, GM, GAFS, CC, BL, L-ATT, ZM, SMA, and HM oversaw the collection of modern worm samples. BLF and MBer provided expertise and advice throughout the study. CMOK, PN, MJS, and SRD analysed and interpreted results. SRD led and performed the bioinformatics analyses, and drafted the manuscript and figures. All authors have read and approved the final version of the manuscript.

## Methods

### Sampling of ancient and modern *Trichuris* spp. from humans and animals

Sampling details specific to each group or population of parasites are outlined below. As the focus of the study is on the genetics of whipworms, no patient-specific data or identifiers were included in any analyses performed. In all cases where parasites were collected from human patients, informed written consent was obtained from all participants or their guardians (in cases where the participant was a child) after being informed about the study in English and/or the local language. Samples were collected in line with ethical consideration and clearances as follows:

#### Ancient

*Trichuris* sp. eggs were isolated from ancient latrine samples in which a metagenomic study has previously identified DNA from *T. trichiura* ^13^. The eggs were isolated from ancient human latrine or deposit sites from Denmark (1000—1700 AD), the Netherlands (1350—1850 AD), and Lithuania (1550—1580 AD) (**Supplementary Fig. 1**).

#### China

*T. trichiura* samples, previously defined as genetically distinct from pig-derived *Trichuris* (i.e. *T. suis*) by mitochondrial genome and nuclear genetic markers ^52^, were collected directly from the caecum of a human patient during surgery in Zhanjiang People’s Hospital in Zhanjiang, Guangdong Province, China (CHN_GUA). These worms were sent to the Department of Parasitology, Guangdong Medical University, for identification.

#### Colobus guereza kikuyensis

Worms, previously defined as *Trichuris colobae n. Sp*. ^53^, were collected from the caecum during necropsy of *Colobus guereza kikuyensis* (Mantled Guereza or Eastern Black-and-White Colobus) (ESP_MAL [zoo]), which died of natural causes at the Fuengirola Zoo (Málaga, Spain).

#### Papio hamadryas

Worms from baboons were recovered from the caecum during routine post mortem examination of animals culled for management reasons at Copenhagen Zoo, Denmark ^23^.

#### Honduras

Samples were collected as a joint effort by Ana Sanchez (Brock University, Canada), Gustavo A. Fontecha and Gabriela Matamoros (Universidad Nacional Autónoma de Honduras (UNAH), Honduras) (HND_OLA and HND_SAL). Sample collection protocols were reviewed and approved by the Brock University Bioscience Research Ethics Board and Comité de Ética de Investigación – Maestría en Enfermedades Infecciosas y Zoonóticas – Facultad de Ciencias – UNAH.

#### Trachypithecus francoisi

*Trichuris* spp. samples were collected during necropsy from the caecum of a captive *Trachypithecus francoisi* (François Leaf-Monkey), which was humanely euthanised due to acute gastric dilation (People’s Republic of China) (CHN_GUA [zoo]) ^54^.

#### Uganda

Worms were recovered after anthelmintic treatment from the faeces of children in Uganda (UGA_KAB) as previously described ^55^. Permission was obtained from the Ministry of Health, and the National Council of Science and Technology in Uganda and the Danish Central Medical Ethics Committee recommended the study. A subset of worms was recovered after anthelmintic treatment from a Danish patient infected with *T. trichiura* from Uganda (UGA_DNK) as previously described ^3^.

#### Cameroon

Stool samples were collected with a focus on monitoring the efficacy of mebendazole for the treatment of soil-transmitted helminths ^6^. Permission was obtained from the National Ethics Committee in Yaoundé, Cameroon (Sep 2011).

#### Tanzania

Stool samples were collected as previously described ^6^. Permission was obtained from the Ministry of Health, Zanzibar Revolutionary Government, Tanzania (July 2012).

Individual worms were washed extensively in tap water or saline solution and preserved in 70% ethanol. When possible, the sex of individual worms was determined through microscopic examination before DNA extraction.

### DNA extraction and high-throughput sequencing

Ancient samples were processed for egg extraction in a dedicated paleoparasitological laboratory at the Department of Plant and Environmental Sciences (PLEN), University of Copenhagen. Helminth eggs were concentrated by selecting particles based on flotation in high-density liquid sedimentation and size filtration. The egg samples were then processed for DNA extraction in dedicated aDNA laboratories at the Centre for Geogenetics (CGG), University of Copenhagen, in accordance with strict aDNA-specific requirements. DNA extraction was performed using the PowerLyzer PowerSoil DNA isolation kit (MO BIO Laboratories, Carlsbad, California) with minor modifications. The complete protocol has been described previously ^13^.

DNA from modern worm samples were prepared as follows: (i) for samples collected from stool following treatment with anthelmintics, directly from a human patient via colonoscopy, and from baboon, DNA was extracted from whole adult worms using the MasterPure^™^ DNA Purification kit (Epicentre Biotechnologies) at the Department of Veterinary Disease Biology (VDB), UCPH following the manufacturer’s protocol with the following exception: worms were first homogenised in 300 μL lysis solution using a matching disposable plastic pestle and then incubated at 56°C for three hours; (ii) for the colobus samples, DNA extraction from whole adult worms was performed at the Departamento de Microbiología y Parasitología, Universidad de Sevilla, Sevilla, Spain, using a DNeasy Blood & Tissue Kit (Qiagen) according to the manufacturer’s protocol; (iii) for the leaf monkey samples, DNA extraction was performed at the Department of Parasitology, Lanzhou Veterinary Research Institute, Lanzhou, China as previously described ^54^. DNA (400 ng) was fragmented to approximately 300—700 bp using a BioRuptor (Diagenode) using 2 or 4 cycles of 15 sec ON / 90 sec OFF at instrument setting HIGH. Fragmented DNA was concentrated using a MinElute kit (Qiagen), after which DNA concentration was determined (Qubit 2.0 dsDNA HS kit), and library size confirmed (Agilent Bioanalyzer High Sensitivity DNA kit).

DNA sequencing libraries were prepared for all sample types, modern and ancient DNA extracts, using the NEBNext DNA Sample Prep Master Mix Set for 454 (E6070) kit with a modified protocol previously described for ancient samples ^13^. DNA libraries were amplified in a single PCR reaction of 50 μL using: half of the prepared DNA library (12.5 μL), 5 U Taq Gold (Life Technologies), 1x buffer Gold, 2 mM MgCl_2_, 0.25 mM dXTP, 0.2 mM of primers (PE1.0 and Illumina multiplex primer), for 8—16 cycles depending on library strength. Libraries were sequenced using 100 bp single-end (ancient samples) or 100 bp paired-end (modern samples) chemistry on a HiSeq 2000/2500 platform at The Danish National High-Throughput DNA Sequencing Centre (now Centre for GeoGenetics Sequencing Core).

### Raw read processing and mapping to the reference genome

Raw reads were first processed using AdapterRemoval2 ^56^; for both SE and PE reads, adapters were removed and N bases trimmed, and for PE reads, R1 and R2 reads were collapsed where possible. Where multiple lanes of data were generated, trimmed read sets were merged before mapping.

Mapping was performed using BWA-MEM ^57^ to an unpublished but significantly improved reference genome of *T. trichiura* (available here: ftp://ngs.sanger.ac.uk/production/pathogens/parasites/Trichuris/trichuria/). Originally described by Foth *et al*. ^58^, the new assembly used here is larger (80.57 Mb vs 75.49 Mb), more contiguous (N50 = 11.3 Mb [vs 0.07 Mb] & N50n = 2 [vs 265]), and in fewer scaffolds overall (n = 113 vs 4156) relative to the published version. For the ancient samples, trimmed SE reads were mapped, whereas for the modern samples, trimmed PE and SE reads (merged PE and trimmed SE) were mapped independently and subsequently merged. For all samples, duplicate reads were identified using picard MarkDuplicates (http://broadinstitute.github.io/picard/).

Putative deamination damage (represented by excessive C-to-T and G-to-A substitutions at the ends of reads) was assessed using PMDtools (https://github.com/pontussk/PMDtools) ^59^. This analysis revealed bias in the terminal 2 bp, particularly in the ancient samples; to account for this, the mapped reads were trimmed by 2 bp for all samples (ancient and modern) using bamUtils trimBam ^60^. Genome coverage was determined using samtools bedcov per scaffold and in 100 kb windows. To explore putative causes of mapping variation, we also ran Kraken2 ^61^ to estimate the degree of contamination in the raw sequencing reads. This analysis showed that while each sample contained a small degree of “contamination” evidenced by hits to the kraken database (minikraken2_v1_8GB), it did not explain overall variation in mapping. Quality control and quantitative comparison between samples was undertaken at each stage of the pipeline and visualised using MultiQC ^62^.

### Variant calling

Variant calling was performed with GATK Haplotype Caller, first by generating per sample GVCF files (--min-base-quality-score 20 --minimum-mapping-quality 30 --standard-min-confidence-threshold-for-calling 30), followed by joint genotyping of the sample cohort. Heterozygosity was adjusted from default settings based on estimates from raw reads using GenomeScope ^63^. To improve efficiency, each task was split by sample and by scaffold and run in parallel before merging to produce the final call sets.

The raw output was split into mitochondrial and nuclear variants and then split again into single nucleotide variants (SNV) and indels and filtered independently. A “hard filtering” approach was employed based on removing relevant tails of variant distributions (typically the upper and/or lower 1%) on the following quality metrics: QUAL, DP, MQ, SOR, FS, QD, MQRankSum, and ReadPosRankSum. Variants were further filtered to ensure they met the following criteria: minimum and maximum alleles = 2; minor allele > 0.02; Hardy Weinberg Equilibrium = >1E-6 (nuclear variants); per sample missingness > 0.8; per genotype depth > 3. Finally, variants were removed if they were found in regions of the genome where the reference genome-guided mapping of kmers (k = 35) was poor or not unique determined using SNPable (http://lh3lh3.users.sourceforge.net/snpable.shtml), retaining regions of the genome where overlapping 35-mers mapped uniquely and without 1-mismatch (75.02% of the genome was retained).

### Population structure analyses

Broad-scale genetic relatedness between samples and populations was explored by principal component analysis using the R package SNPrelate ^64^. These analyses were performed on both mitochondrial and nuclear variants separately and with and without the colobus and leaf monkey samples, which were found as outliers in all analyses.

To further examine the relationships between samples we generated identify-by-state (IBS) and IBS covariance matrices using ANGSD ^27^ (parameters: -minMapQ 30 -minQ 20 -GL 2 - doMajorMinor 1 -doMaf 1 -SNP_pval 2e-6 -doIBS 1 -doCounts 1 -doCov 1 -makeMatrix 1 - minMaf 0.05). This approach took mapped reads in a bam file directly as input (as opposed to variants in a VCF), and calculated genotype likelihoods per variant site, which are more sensitive to poorer quality and low coverage data.

### Phylogenetics of reconstructed and publicly available mitochondrial genomes

Preprocessed sequence data from modern samples were mapped to published mitochondrial genomes of *T. trichiura* (accession numbers: KT449826, GU385218 and HG806815 [an extract from the *T. trichiura* whole-genome shotgun sequence]), *T. suis* (KT449822, KT449823, GU070737), *T. discolor* (JQ996231), *T. ovis* (JQ996232) and that of the Leaf Monkey, *Trichuris* sp. (KC461179) reporting only the best hit. The published genome with the most hits was then used as a reference in guided assemblies using the mitochondrial baiting and iterative mapping approach, MITObim (v1.8) ^65^. Subsequent manual curation of the assemblies was performed to remove redundant, overlapping sequence at the ends of the mitochondrial contigs, and the starting position was adjusted to the first codon of the COXI gene. For the ancient samples, consensus mitochondrial genome sequences have previously been called for ten samples (of 13 in total) that provided 100% bases with coverage to the KT449826 *T. trichiura* reference sequence ^13^. Consensus sequences were annotated using the MITOS Webserver ^66^. Assembled mitochondrial genomes were aligned together with all *Trichuris* spp. reference genomes publicly available at NCBI using mafft v7 ^67^, with the global-pair setting at 1,000 iterations. A neighbour-joining tree was built using the Jukes-Cantor nucleotide distance model and 1,000 bootstraps in CLC Sequence Viewer 7 (Qiagen).

### Admixture and population demographic analyses

Admixture was determined using NGSadmix ^68^. Briefly, genotype likelihoods were extracted using vcftools version 0.1.16 (--max-missing 1 --BEAGLE-PL), after which NGSadmix (-P 4 - minMaf 0.05 -misTol 0.9) was run over a range of K values from 2 to 10. The optimal value of K was determined by iteratively running NGSadmix as above five times, changing the seed value on each run. The log-likelihoods of each run (all iterations of K = 2—10, s = 1—5) were used to determine the optimal value of K using Evanno’s method ^69^ on the Clumpak webserver ^70^.

Treemix ^71^ was performed on the nuclear dataset, including leaf and colobus monkey samples as outliers. Variants were first filtered using vcftools (--max-missing 1), followed by further pruning to minimise variants in linkage disequilibrium. Customs scripts (ldPruning.sh & vcf2treemix.sh) were modified from https://github.com/speciationgenomics/scripts. The optimal number of migration edges was estimated using the R package OptM (https://cran.r-project.org/web/packages/OptM/).

*f*-statistics were calculated using ADMIXTOOLS ^72^ using the *qp3Pop* tool to calculate *f*3 data. All combinations of source 1 and source 2 populations were determined, using either baboon or Honduras samples as the outgroup. Customs scripts (convertVCFtoEigenstrat.sh) were modified from https://github.com/joanam/scripts. Standard error was calculated using a weighted block jackknife, using three blocks. The Z score was determined for each comparison to test the deviation from 0 (no allele sharing); a Z score of three or greater was deemed significant.

Population demographics were determined using SMC++ ^73^. For each population, variant sites present in all individuals were extracted using vcftools version 0.1.16 (--max-missing 1), after which smc++ vcf2smc was run per scaffold. Estimated population sizes were fit to the data for all scaffolds using smc++ estimate, which used the nematode *C. elegans* mutation rate of 2.9e-9 mutations per site per generation ^74^ as a proxy for *T. trichiura* mutation rate, which is currently unknown. Finally, the effective population size per generation was scaled based on an estimated generation time; while unknown precisely, the prepatent period has been estimated to range between 13 to 16 weeks ^3^ and, therefore, we chose a time period of three months or four generations per year. As the ancient samples were a pooled population of eggs rather than individual parasites, they violated assumptions of the model and were, therefore, removed from the analysis.

Kinship between samples within populations was assessed using vcftools (--relatedness2) and NGSRelate ^75^.

### Genome-wide genetic diversity analyses

Genome-wide nucleotide diversity, Tajima’s *D*, and pairwise *F*_ST_ was determined using vcftools in 50 kb non-overlapping sliding windows. To improve the visualisation of the genome-wide comparisons, analyses were restricted to scaffolds in chromosomal linkage groups, which represented 89.83% of the whole genome assembly.

The annotation of the unpublished genome assembly used in this analysis was not available at the time of analysis. To enable the identification of genes within outlier regions of the genome-wide analyses, we performed a liftover of existing gene models and gene identifiers from the published version of the *T. trichiura* assembly currently available in WormBase Parasite ^76^ (https://parasite.wormbase.org/Trichuris_trichiura_prjeb535/Info/Index/, Version: WBPS15) using liftoff ^77^. By retaining gene identifiers from the original assembly, cross-validation of gene hits within WormBase ParaSite could be performed. In total, 8451 of 9650 gene features (∼88%) were transferred; of the genes that did not transfer, ∼80% were classified as contamination based on hits to the uniprot_reference_proteome database using DIAMOND ^78^. Gene ontology term analysis was performed using gProfiler ^77,79^, applying a g:SCS multiple testing correction method and a significance threshold of 0.05, restricting the complete geneset to only genes that had successfully been transferred from the original annotation.

Analysis of variation in β-tubulin was performed using vcftools (--site-pi --maf 0.01), using a bed file of exon coordinates derived from manual curation of the gene.

## Data availability

Raw sequencing data is available under the European Nucleotide Archive (ENA) study accession ERP128004. The *Trichuris trichiura* genome assembly is available at ftp://ngs.sanger.ac.uk/production/pathogens/parasites/Trichuris/trichuria/trichuris_trichiura.fa.

## Code availability

Custom code to analyse data and reproduce the figures presented is available at https://stephenrdoyle.github.io/ancient_trichuris/.

