## Supplementary Figure for "Population genomics of ancient and modern *Trichuris trichiura*"

**List of Figures**

- **Supplementary Figure 1.** The estimated age of sampling sites from which ancient *Trichuris trichiura* samples were collected.
- **Supplementary Figure 2.** Evidence of deamination in raw sequencing reads.
- **Supplementary Figure 3.** Relative genome-wide sequencing coverage of all samples sequenced.
- **Supplementary Figure 4.** Identity by state (IBS) between all samples based on nuclear markers.
- **Supplementary Figure 5.** Covariance of IBS between all samples based on nuclear markers.
- **Supplementary Figure 6.** Genetic analysis of leaf monkey and colobus monkey isolates.
- **Supplementary Figure 7.** Extended admixture analysis.
- **Supplementary Figure 8.** Extended treemix analysis.
- **Supplementary Figure 9.** Comparison of private and shared variation between Ugandan, Chinese, and American isolates.
- **Supplementary Figure 10.** Comparison of nucleotide diversity between populations.
- **Supplementary Figure 11.** Comparison of Tajima's D between populations.
- **Supplementary Figure 12.** Analysis of variation within and surrounding the β-tubulin gene.

**List of Tables (see accompanying document)**

- **Supplementary Table 1.** Sample metadata and sequencing accession numbers
- **Supplementary Table 2.** Genome mapping data by sample, including nuclear and mitochondrial genome coverage, and deamination statistics.
- **Supplementary Table 3.** Description of genes in regions of high genetic differentiation between samples from China and Uganda
- **Supplementary Table 4.** Description of genes in regions of high genetic differentiation between samples from Uganda and the Americas

- **Supplementary Table 5.** Description of genes in regions of high genetic differentiation between samples from China and the Americas
- **Supplementary Table 6.** Description of genes in regions of high genetic differentiation between samples from Baboons and Uganda.

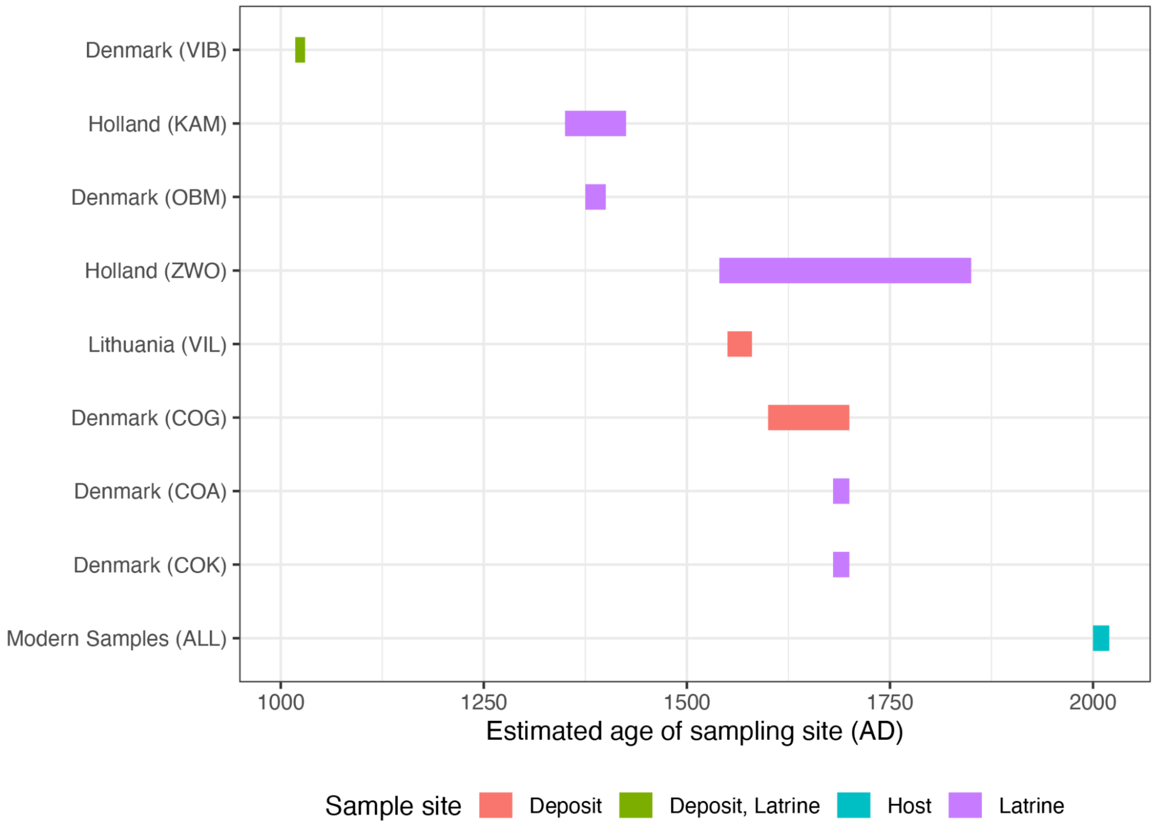


**Supplementary Figure 1. The estimated age of sampling sites from which ancient *Trichuris trichiura* samples were collected.**

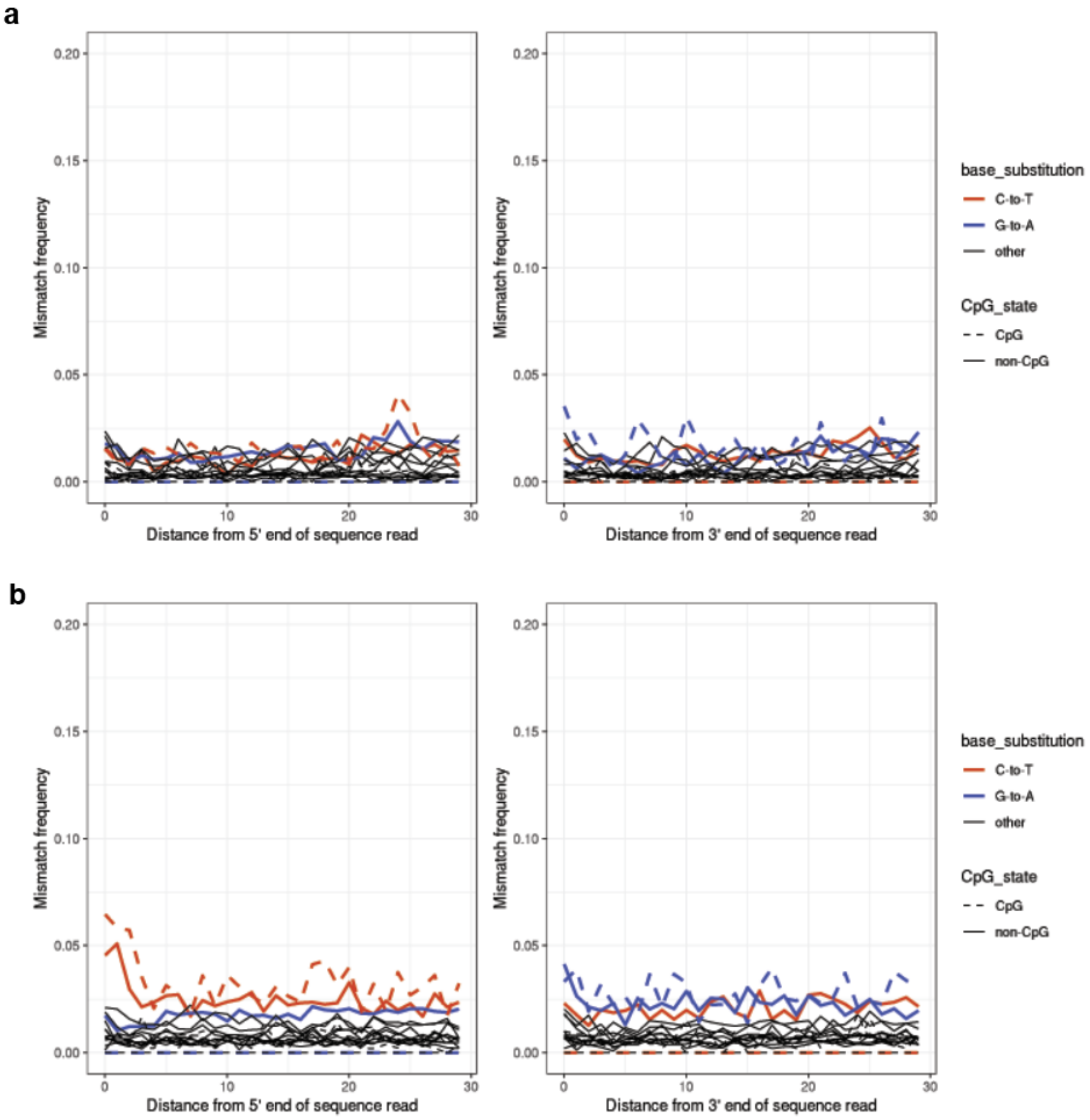


**Supplementary Figure 2. Evidence of deamination in raw sequencing reads.** Deamination of cytosines is a common artefact in ancient DNA, evidenced by an increase in C-to-T and G-to-A substitutions. **a**. Example of a modern sample (MN_CHN_GUA_HS_001). **b**. Example of an ancient sample (AN_DNK_COG_EN_0012), showing an increase in C-toT frequency at the beginning of the reads.

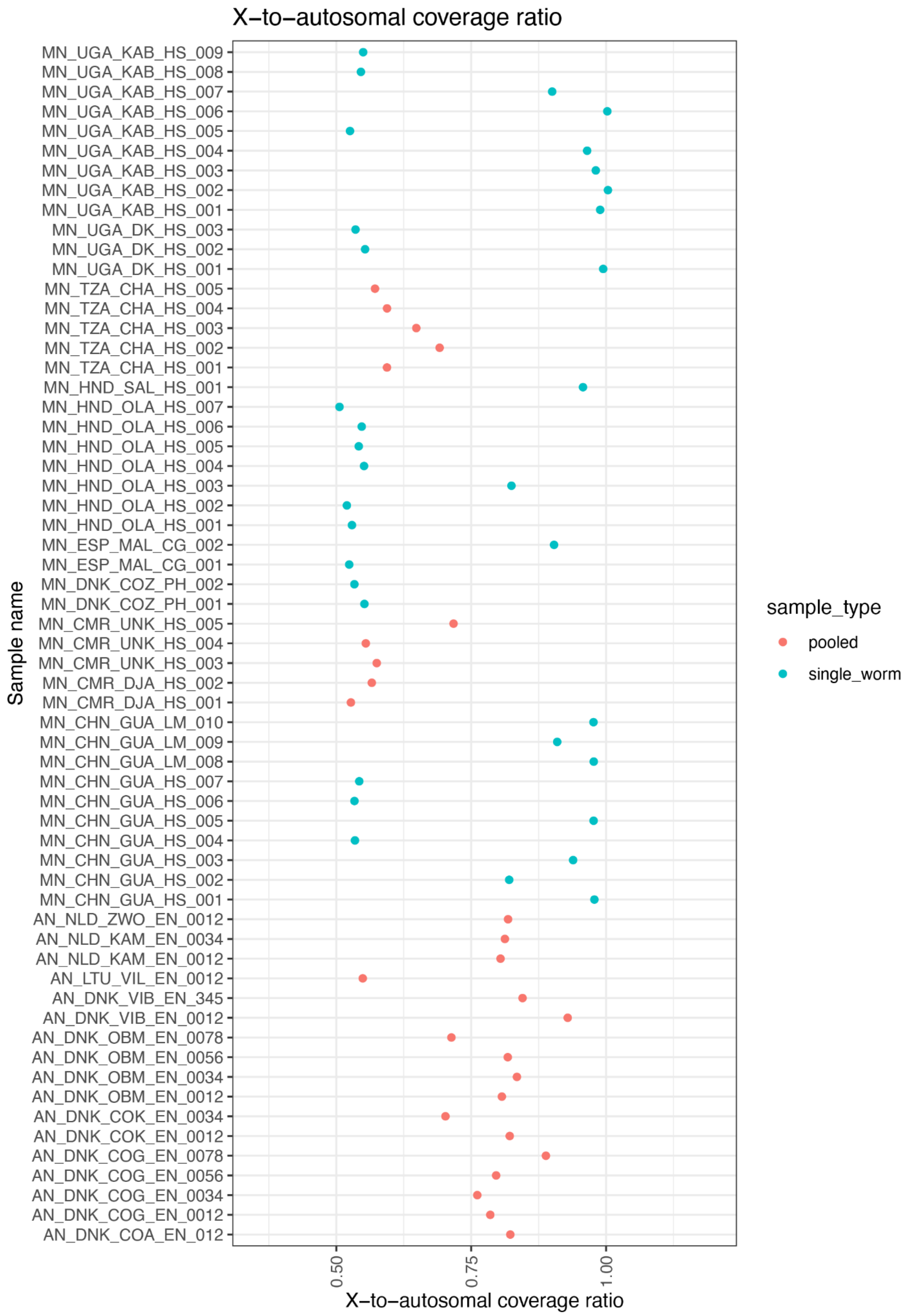

X−to−autosomal coverage ratio
MN_UGA_KAB_HS_009
MN_UGA_KAB_HS_008
MN_UGA_KAB_HS_007
MN_UGA_KAB_HS_006
MN_UGA_KAB_HS_005
MN_UGA_KAB_HS_004
MN_UGA_KAB_HS_003
MN_UGA_KAB_HS_002
MN_UGA_KAB_HS_001
MN_UGA_DK_HS_003
MN_UGA_DK_HS_002
MN_UGA_DK_HS_001
MN_TZA_CHA_HS_005
MN_TZA_CHA_HS_004
MN_TZA_CHA_HS_003
MN_TZA_CHA_HS_002
MN_TZA_CHA_HS_001
MN_HND_SAL_HS_001
MN_HND_OLA_HS_007
MN_HND_OLA_HS_006
MN_HND_OLA_HS_005
MN_HND_OLA_HS_004
MN_HND_OLA_HS_003
MN_HND_OLA_HS_002
MN_HND_OLA_HS_001
MN_ESP_MAL_CG_002
MN_ESP_MAL_CG_001
MN_DNK_COZ_PH_002
MN_DNK_COZ_PH_001
MN_CMR_UNK_HS_005
MN_CMR_UNK_HS_004
MN_CMR_UNK_HS_003
MN_CMR_DJA_HS_002
MN_CMR_DJA_HS_001
MN_CHN_GUA_LM_010
MN_CHN_GUA_LM_009
MN_CHN_GUA_LM_008
MN_CHN_GUA_HS_007
MN_CHN_GUA_HS_006
MN_CHN_GUA_HS_005
MN_CHN_GUA_HS_004
MN_CHN_GUA_HS_003
MN_CHN_GUA_HS_002
MN_CHN_GUA_HS_001
AN_NLD_ZWO_EN_0012
AN_NLD_KAM_EN_0034
AN_NLD_KAM_EN_0012
AN_LTU_VIL_EN_0012
AN_DNK_VIB_EN_345
AN_DNK_VIB_EN_0012
AN_DNK_OBM_EN_0078
AN_DNK_OBM_EN_0056
AN_DNK_OBM_EN_0034
AN_DNK_OBM_EN_0012
AN_DNK_COK_EN_0034
AN_DNK_COK_EN_0012
AN_DNK_COG_EN_0078
AN_DNK_COG_EN_0056
AN_DNK_COG_EN_0034
AN_DNK_COG_EN_0012
AN_DNK_COA_EN_012
Sample name
0.50
0.75
1.00
X−to−autosomal coverage ratio
sample_type
pooled
single_worm

**Supplementary Figure 3. Parasite sex determination based on relative X-to-autosome sequencing coverage.** The mean coverage of the X-linked scaffolds (linkage group indicated by "Trichuris_trichiura_1" in reference) was compared with the mean coverage from the autosomal scaffolds (linkage groups "Trichuris_trichiura_2" and "Trichuris_trichiura_3" in reference). Male worm (XY) sex-linked scaffolds present at half-coverage of the autosomes (i.e., ratio = ~0.5), whereas female worms (XX) show equal coverage between sex-linked and autosomal scaffolds (i.e., ratio = ~1). Note that the ancient samples, as well as a subset of the African samples (Tanzania & Cameroon), are pools of eggs and so the ratio of male to females may vary per sample; if an equal sex ratio was present, we would expect a sex/autosome ratio of approximately 0.75.

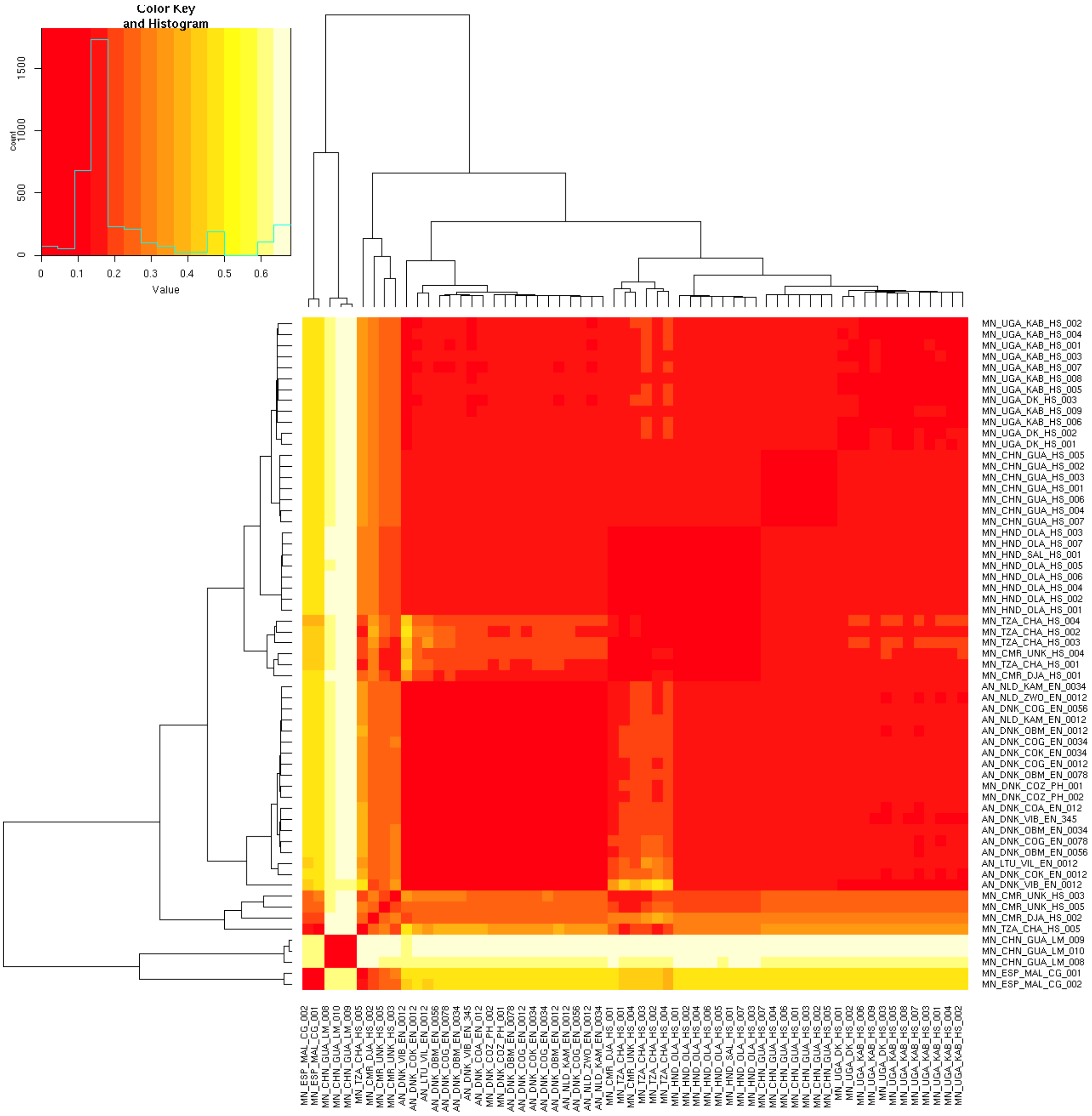


**Supplementary Figure 4. Identity by state (IBS) between all samples based on nuclear markers.** Heatmap shows IBS between pairs of samples calculated using ANGSD; red (values approaching zero) indicates a high degree of shared alleles, whereas yellow (values approaching one) represents a low degree of shared alleles. All samples, including low coverage African and divergent animals, are shown.

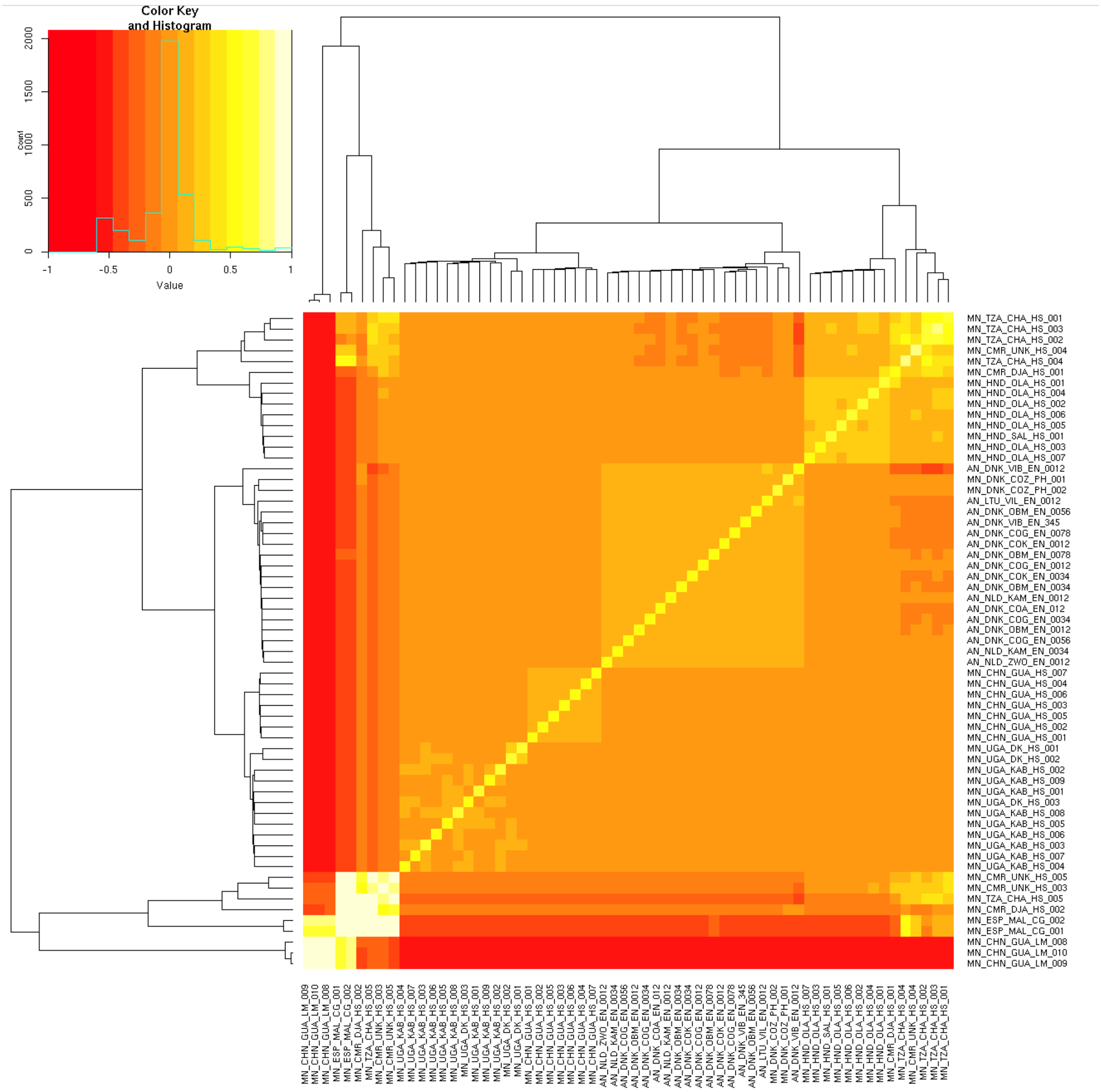


**Supplementary Figure 5. Covariance of IBS between all samples based on nuclear markers.** Heatmap shows covariance in IBS between pairs of samples calculated using ANGSD; red (values approaching negative one) indicates poor covariance between samples, whereas yellow (values approaching positive one) represents strong/high covariance between samples. All samples, including low coverage African and divergent animal samples, are shown.

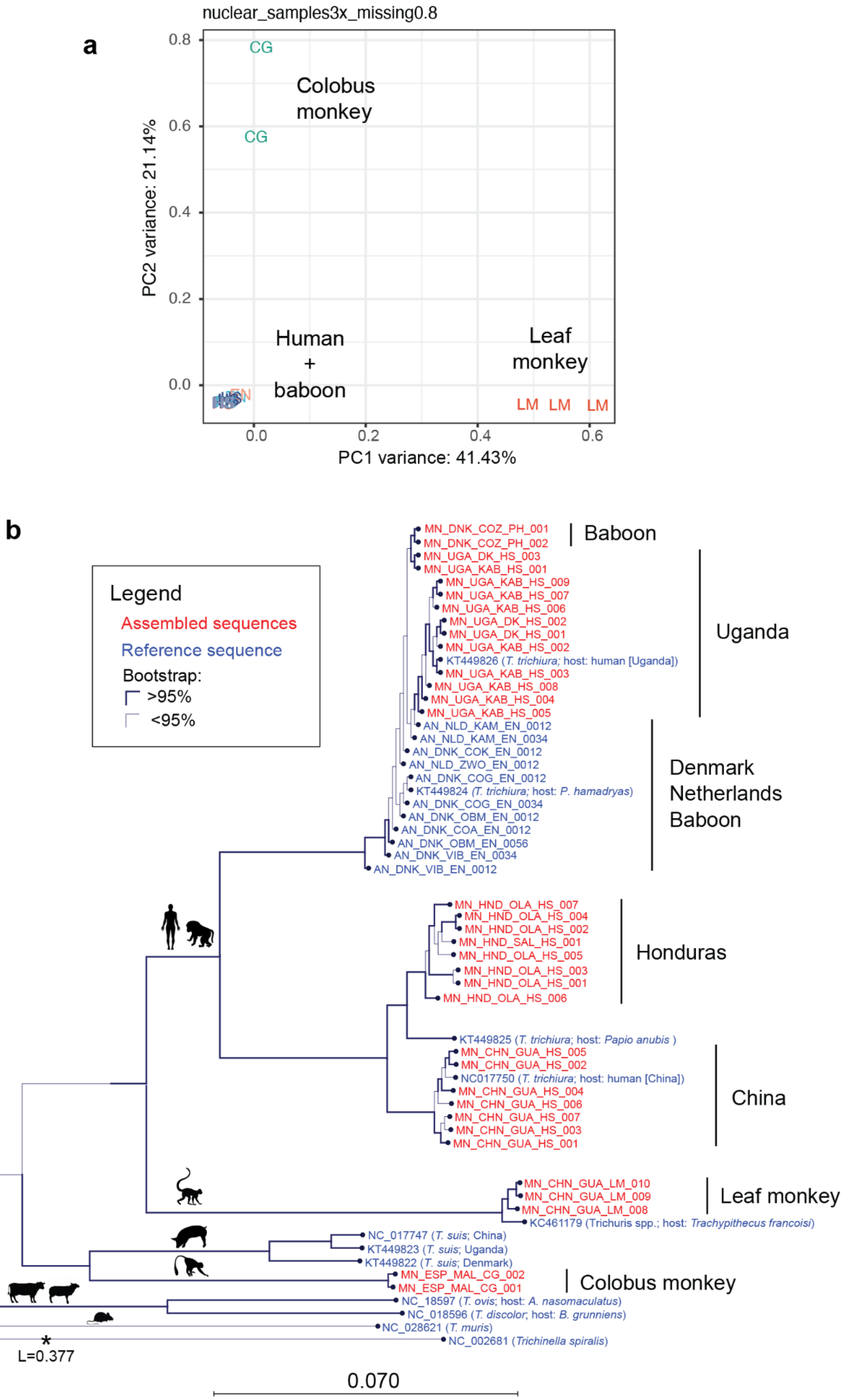
a
nuclear_samples3x_missing0.8
CG
CG
Colobus monkey
Human + baboon
Leaf monkey
LM LM LM
PC2 variance: 21.14%
PC1 variance: 41.43%
0.8
0.6
0.4
0.2
0.0
0.0
0.2
0.4
0.6
b
Legend
Assembled sequences
Reference sequence
Bootstrap:
>95%
<95%
MN_DNK_COZ_PH_001
MN_DNK_COZ_PH_002
MN_UGA_DK_HS_003
MN_UGA_KAB_HS_001
MN_UGA_KAB_HS_009
MN_UGA_KAB_HS_007
MN_UGA_KAB_HS_006
MN_UGA_DK_HS_002
MN_UGA_DK_HS_001
MN_UGA_KAB_HS_002
KT449826 (T. trichiura; host: human [Uganda])
MN_UGA_KAB_HS_003
MN_UGA_KAB_HS_008
MN_UGA_KAB_HS_004
MN_UGA_KAB_HS_005
AN_NLD_KAM_EN_0012
AN_NLD_KAM_EN_0034
AN_DNK_COK_EN_0012
AN_NLD_ZWO_EN_0012
AN_DNK_COG_EN_0012
KT449824 (T. trichiura; host: P. hamadryas)
AN_DNK_COG_EN_0034
AN_DNK_OBM_EN_0012
AN_DNK_COA_EN_0012
AN_DNK_OBM_EN_0056
AN_DNK_VIB_EN_0034
AN_DNK_VIB_EN_0012
Baboon
Uganda
Denmark Netherlands Baboon
MN_HND_OLA_HS_007
MN_HND_OLA_HS_004
MN_HND_OLA_HS_002
MN_HND_SAL_HS_001
MN_HND_OLA_HS_005
MN_HND_OLA_HS_003
MN_HND_OLA_HS_001
MN_HND_OLA_HS_006
Honduras
KT449825 (T. trichiura; host: Papio anubis )
MN_CHN_GUA_HS_005
MN_CHN_GUA_HS_002
NC017750 (T. trichiura; host: human [China])
MN_CHN_GUA_HS_004
MN_CHN_GUA_HS_006
MN_CHN_GUA_HS_007
MN_CHN_GUA_HS_003
MN_CHN_GUA_HS_001
China
MN_CHN_GUA_LM_010
MN_CHN_GUA_LM_009
MN_CHN_GUA_LM_008
KC461179 (Trichuris spp.; host: Trachypithecus francoisi)
Leaf monkey
NC_017747 (T. suis; China)
KT449823 (T. suis; Uganda)
KT449822 (T. suis; Denmark)
MN_ESP_MAL_CG_002
MN_ESP_MAL_CG_001
Colobus monkey
NC_18597 (T. ovis; host: A. nasomaculatus)
NC_018596 (T. discolor; host: B. grunniens)
NC_028621 (T. muris)
NC_002681 (Trichinella spiralis)
*
L=0.377
0.070

**Supplementary Figure 6. Genetic analysis of leaf monkey and colobus monkey isolates. a.** PCA of nuclear variants, including highlighting genetically distinct colobus and leaf monkey samples from the closely related, indistinguishable cluster of human and baboon samples. **b.** Neighbour-joining phylogeny of assembled mitochondrial genomes (red) together with publicly available whole mitochondrial genome data (blue; note some of the ancient mitochondrial genomes had previously been described by Soe et al. 2018) for species within the *Trichuris* genus. *Trichinella spiralis* was used as an outgroup. Branch lengths less than 0.002 are set to 0.002.

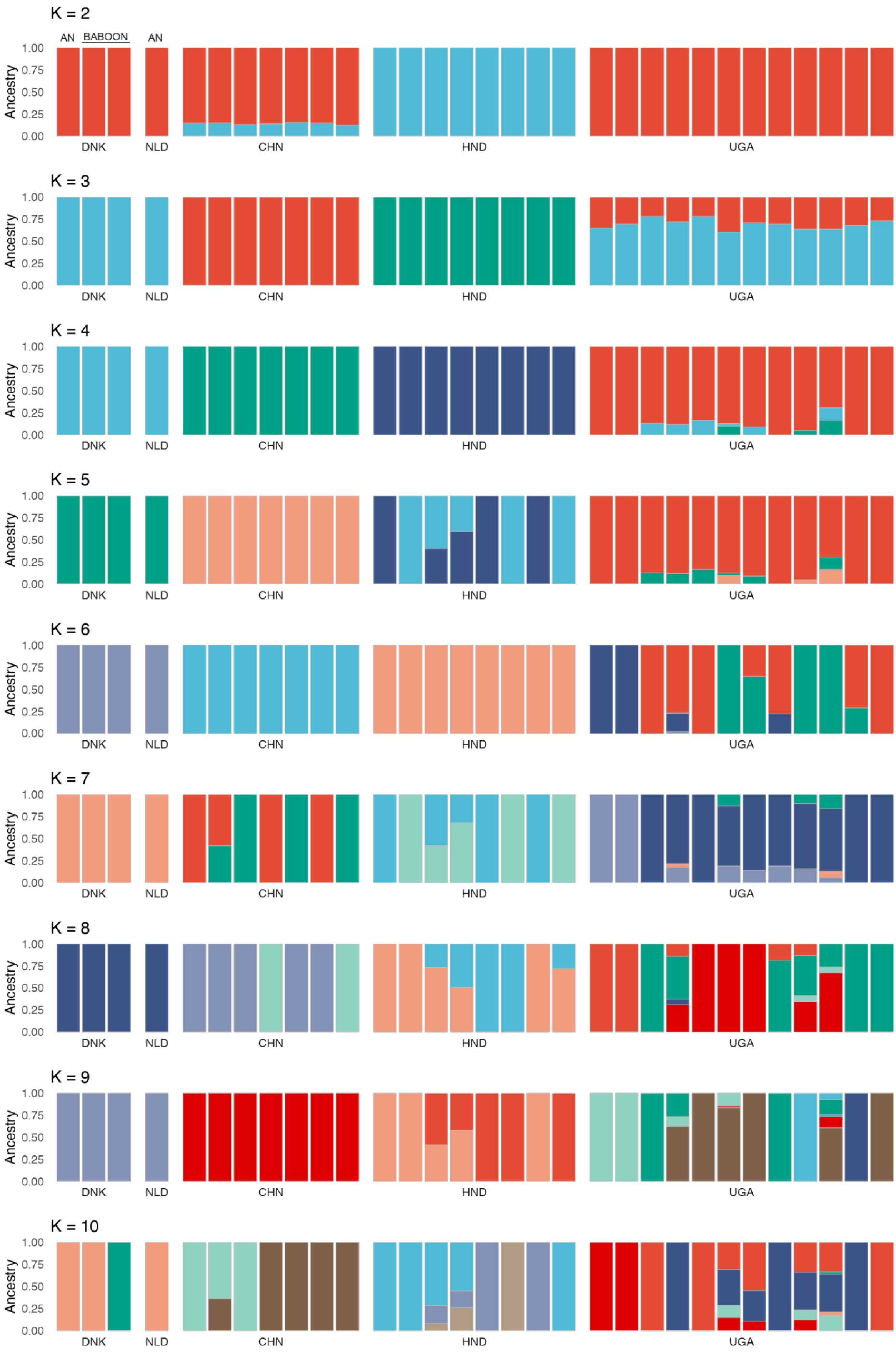
K = 2
AN
BABOON
AN
Ancestry
1.00
0.75
0.50
0.25
0.00
DNK
NLD
CHN
HND
UGA
K = 3
K = 4
K = 5
K = 6
K = 7
K = 8
K = 9
K = 10

**Supplementary Figure 7. Extended admixture analysis.** Analysis of genetic admixture between samples (individual bars; n = 39) and populations (x-axis groups) across a range of theoretical ancestral genetic populations (individual colours) ranging from k = 2 to k = 10 using 857,169 nuclear variants. Ancient samples are represented as the first DNK sample and the NLD sample, and the baboon samples are represented as the second and third DNK samples. The remaining samples are modern from each respective country.

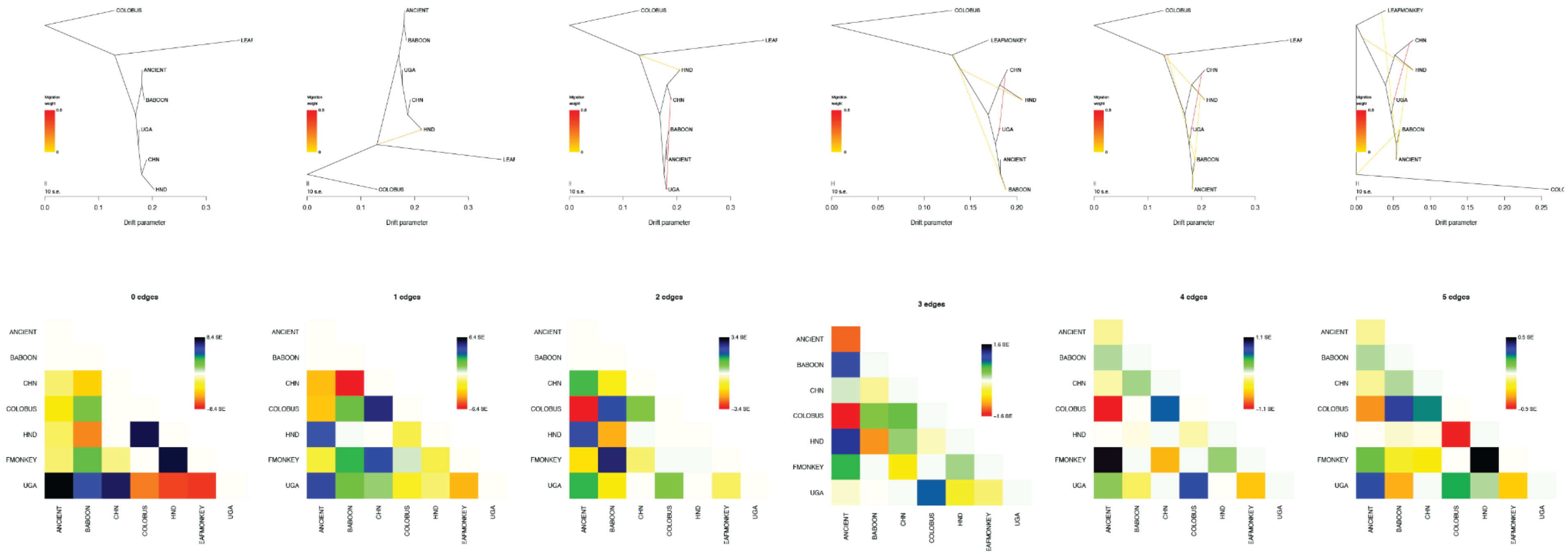

**Supplementary Figure 8. Extended treemix analysis.** Treemix maximum likelihood trees of ancient and modern samples, including Colobus and Leaf monkey samples as outgroups, across a range of hypothetical migration edges that range from no migration events (m = 0 edges) to five migration events (m = 5 edges). The support for those migration edges - the migration weight - is represented by the coloured line joining populations. We also present matrices of the pairwise residuals below each tree. Positive residual values between pairs of populations indicate that they are more closely related to each other than is inferred by the best-fit tree, and therefore, provide a finer-grained view of potential admixture events.

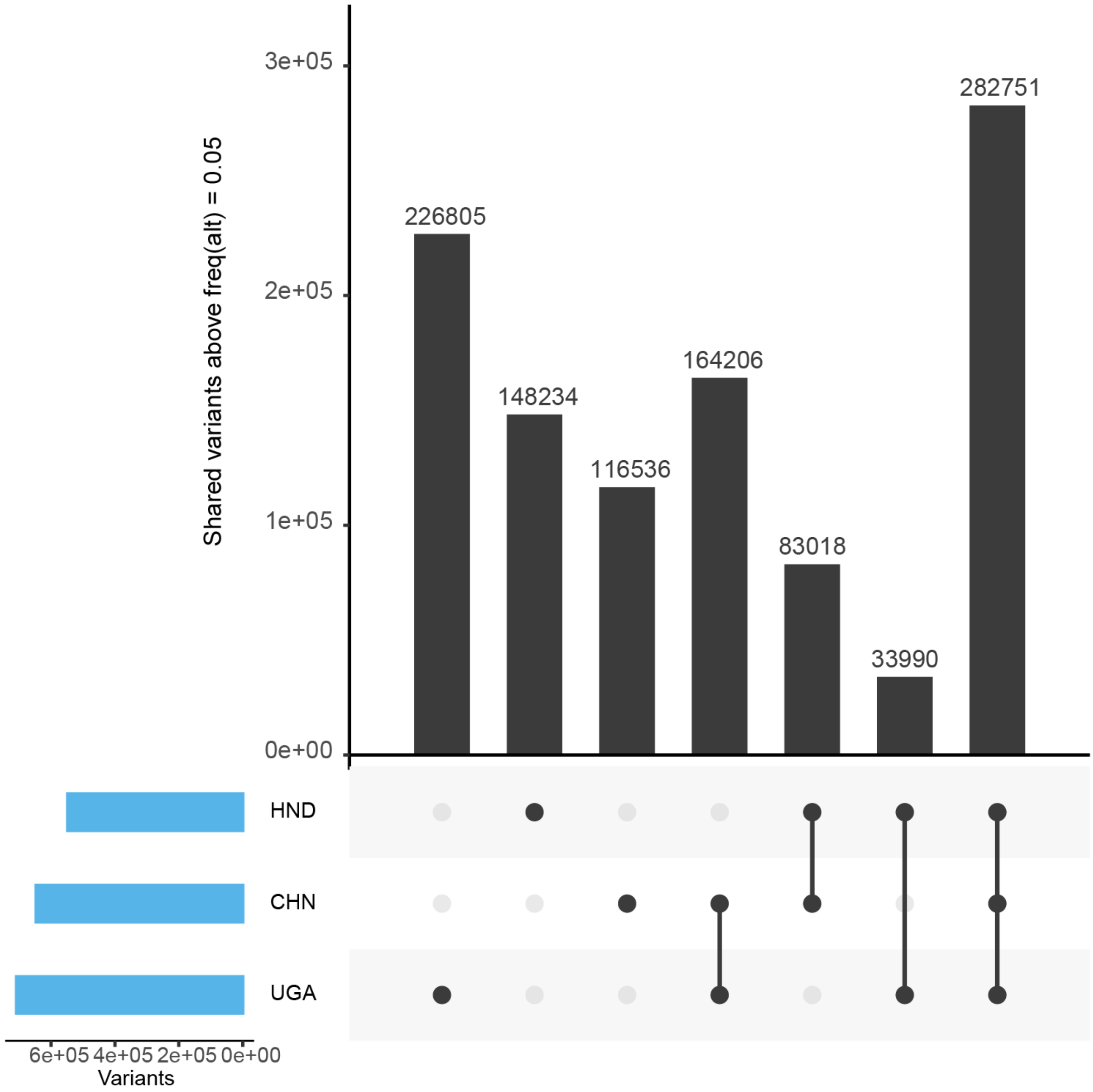


**Supplementary Figure 9. Comparison of private and shared variation between Ugandan, Chinese, and Honduran isolates.** Nuclear variants were subset into groups depending on whether they were presented in a single population (ie. private), shared by only two populations, or by all three populations. Given the step-wise hypothesis by which humans migrated from Africa to Asia and then subsequently to the Americas, the rationale for this analysis was to identify the proportion of variation shared by Ugandan and Honduran populations that is not present in the Chinese population, which may indicate independent translocation.

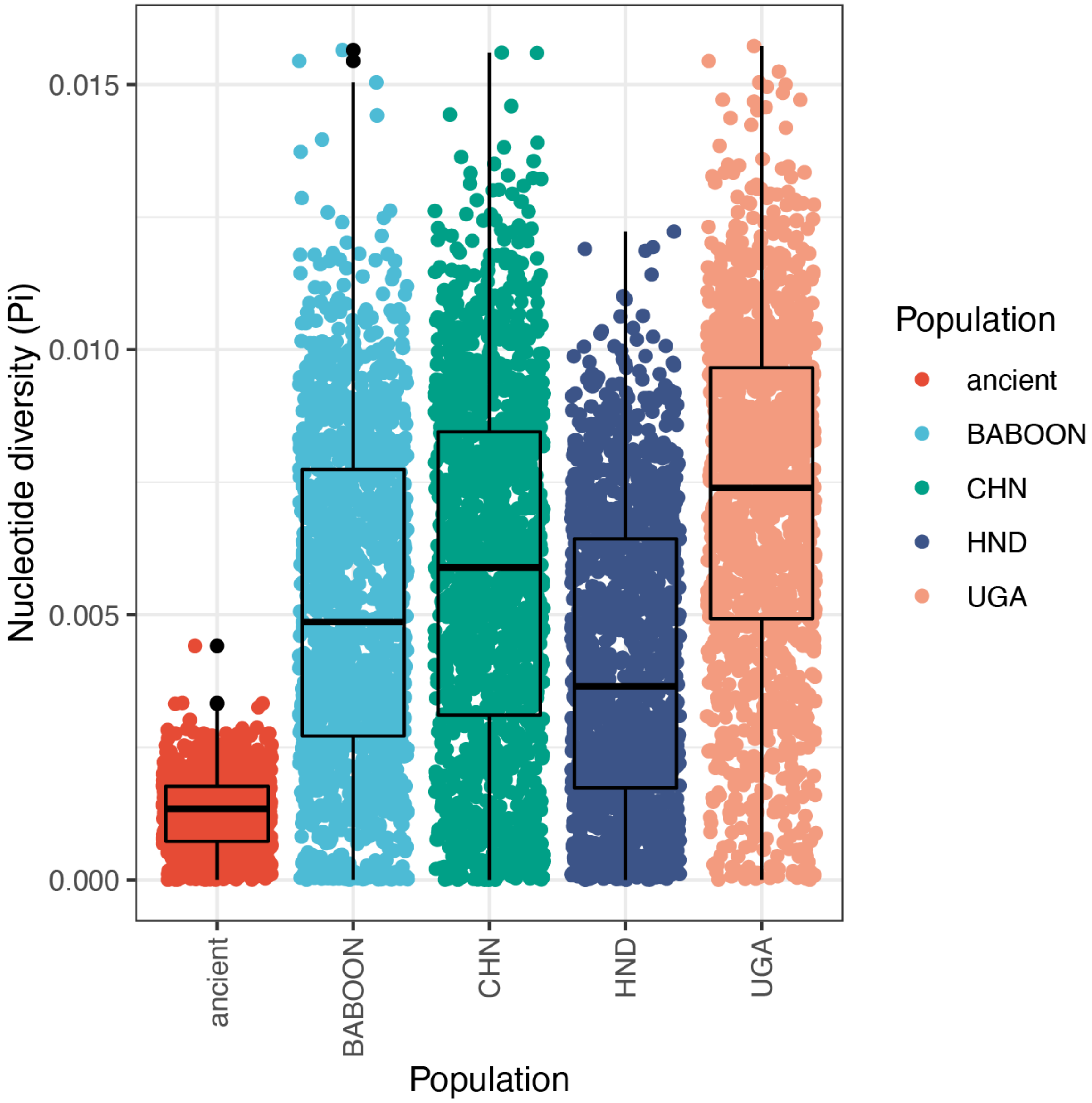


**Supplementary Figure 10. Comparison of nucleotide diversity between populations.** Diversity is measured in 50 kb non-overlapping windows throughout the genome.

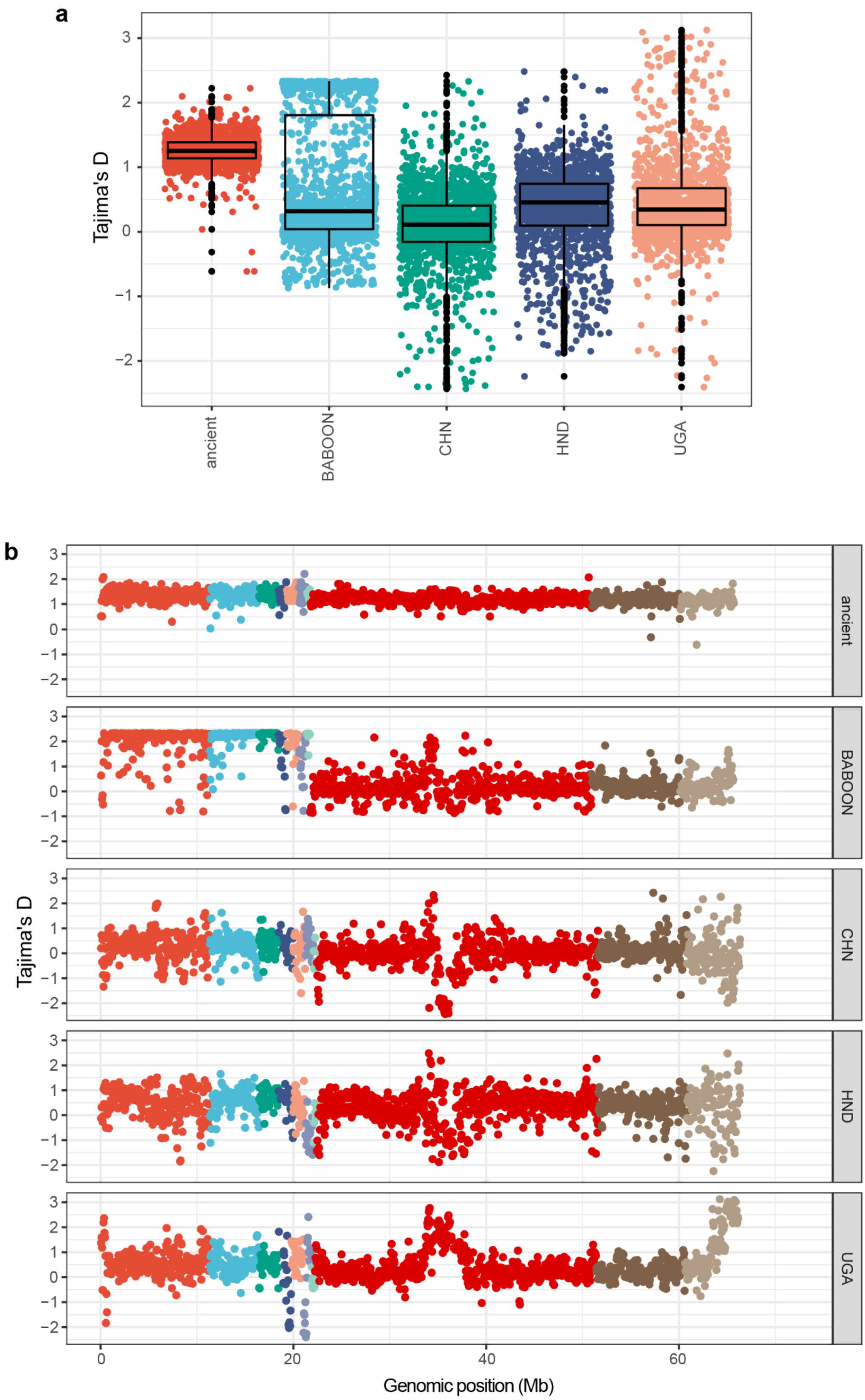


**Supplementary Figure 11. Comparison of Tajima's D between populations.** Tajima's D is measured in 50 kbp non-overlapping windows, summarised by their distribution per population (a) and throughout the genome (b).

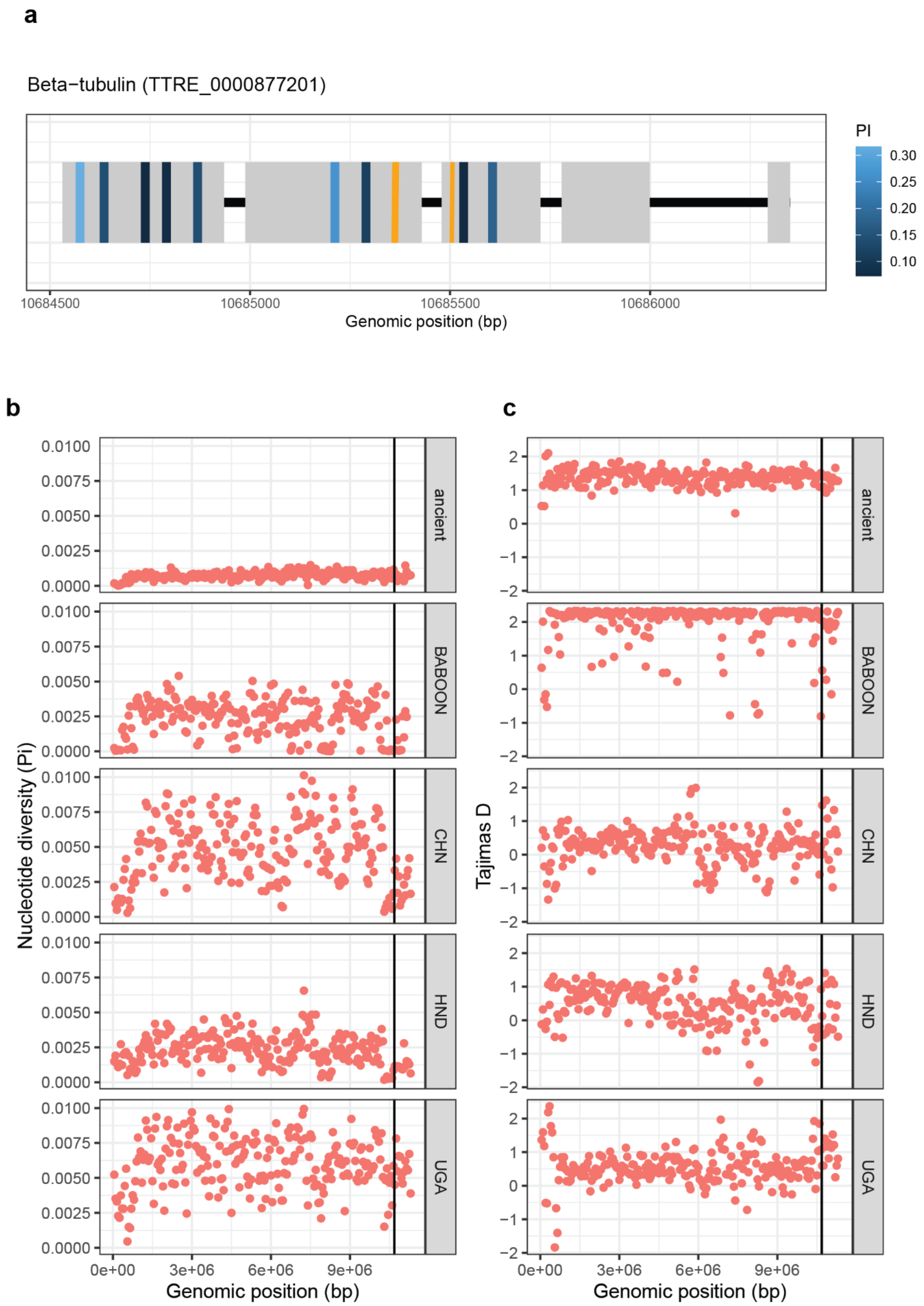
a
Beta−tubulin (TTRE_0000877201)
PI
0.30
0.25
0.20
0.15
0.10
10684500
10685000
10685500
10686000
Genomic position (bp)
b
Nucleotide diversity (Pi)
0.0100
0.0075
0.0050
0.0025
0.0000
ancient
BABOON
CHN
HND
UGA
0e+00
3e+06
6e+06
9e+06
Genomic position (bp)
c
Tajimas D
2
1
0
−1
−2
ancient
BABOON
CHN
HND
UGA
0e+00
3e+06
6e+06
9e+06
Genomic position (bp)

**Supplementary Figure 12. Analysis of variation within and surrounding the β-tubulin gene. a.** Gene structure of the β-tubulin gene (TTRE_0000877201), with exons presented as grey boxes separated by black lines representing introns. Variants identified in the global cohort are indicated by coloured lines: the orange lines represent the positions of canonical resistance-associated variants - P167, P198, and P200 - at which no variation was identified here, whereas the blue lines show variation present, the frequency measured as nucleotide diversity which is indicated by the colour scale shading. **b.** Nucleotide diversity (measured in 50 kbp non-overlapping windows) in the Trichuris_trichiura_1_001 scaffold containing the β-tubulin gene, indicated by the black vertical line, in each population. **c**. Tajima's D (measured in 50 kbp non-overlapping windows) in the Trichuris_trichiura_1_001 scaffold containing the beta-tubulin gene, indicated by the black vertical line, in each population.
